# CD4^+^ T cell senescence is associated with reduced reactogenicity in severe/critical COVID-19

**DOI:** 10.1101/2023.08.04.551565

**Authors:** Jie Zhang, Chun Chang, Zhaoyuan Liang, Tingting Hu, Zhongnan Yin, Ying Liang, Ting Zhang, Yanling Ding, Xianlong Li, Xiaoyan Gai, Xiaoxue Yang, Xin Li, Xixuan Dong, Jiaqi Ren, Yafei Rao, Jun Wang, Jianling Yang, Lixiang Xue, Yongchang Sun

**Author notes:** Corresponding Author, Yongchang Sun, Lixiang Xue,. These authors have contributed equally to this work (Jie Zhang, Chun Chang). These authors are corresponding authors.

## Abstract

**Background:** Aging is a critical risk factor for unfavorable clinical outcomes among COVID-19 patients and may affect vaccine efficacy. However, whether the senescence of T cells impact the progression to severe COVID-19 in the elderly individuals remains unclear.

**Methods:** By using flow cytometry, we analyzed the frequency of senescent T cells (Tsens) in the peripheral blood from 100 elderly patients hospitalized for COVID-19 and compared the difference between mild/moderate and severe/critical illness. We also assessed correlations between the percentage of Tsens and the quantity and quality of spike-specific antibodies by ELISA, neutralizing antibody test kit and Elispot assay respectively, cytokine production profile of COVID-19 reactive T cells as well as plasma soluble factors by cytometric bead array (CBA).

**Results:** We found a significant elevated level of CD4^+^ Tsens in severe/critical disease compared to mild/moderate illness and patients with a higher level of CD4^+^ Tsens (>19.78%) showed a decreased survival rate as compared to those with a lower level (<19.78%), especially in the breakthrough infection. The percentage of CD4^+^ Tsens was negatively correlated with spike-specific antibody titers, neutralization ability and COVID-19 reactive IL-2^+^ CD4^+^ T cells. Additionally, IL-2 producing T cells and plasma levels of IL-2 were positively correlated with antibody levels.

**Conclusion:** Our data illustrated that the percentage of CD4^+^ Tsens in the peripheral blood could act as an efficient biomarker for the capacity of spike-specific antibody production and the prognosis of severe COVID-19, especially in the breakthrough infection. Therefore, restoration of the immune response of CD4^+^ Tsens is one of the key factors to prevent severe illness and improve vaccine efficacy in older adults.

## Introduction

Aging is a critical risk factor for COVID-19 disease severity, clinical outcome and may affect vaccine efficacy. Old individuals showed the highest susceptibility to COVID-19, with higher hospitalization rates, severe illness rates and mortality^1 2^. Results of clinical trials on mRNA and recombinant spike protein vaccines indicated relatively low antibody response and safety in individuals older than 60 years^3 4^. However, some older adults benefit from COVID-19 vaccines^5^, suggesting heterogeneity of anti-viral immunity in the old individuals.

The mechanisms of impaired immune responses after infection or vaccination in old individuals were obscure. Recent studies suggested that the senescence of T cells may dampen humoral and cellular immunity to COVID-19 infection or vaccine. Patients with COVID-19 had increased amounts of CD8^+^ T cells that express CD57, a marker of T cell senescence^6^. Moreover, compared to young adults, older individuals had a reduction of vaccine-induced spike-specific antibody and T cell response, which was negatively correlated with senescent CD8^+^ T cells^7–10^. Even if a number of studies have described such alterations, no comprehensive investigation exists yet in patients aged more than 60 years by far. Therefore, there is an urgent need to assess how and to what extent senescent T cells are responsible for the progression of severe disease and suboptimal vaccine responses observed in older individuals.

In the present study, we described the association between senescent T cells, spike-specific antibody and T cell responses, plasma soluble factors and disease severity as well as clinical outcome in a cohort of COVID-19 patients with an advanced age of more than 80 years. Our results demonstrated that CD4^+^ Tsen, with defect in IL-2 production may impaire quantity and quality spike-specific antibody production, consequently, enabling progression to severe disease. This finding suggests the potential of CD4^+^ Tsen as a biomarker to predict compromised immune responses in the elderly and may be relevant for future vaccine strategies especially for the venerable old populations.

### 1. Study design and human specimens

This study was conducted at the Third Hospital of Peking University (Beijing, China). The inpatients admitted to the hospital from December 23, 2022, to January 19, 2023, who had been confirmed with SARS-CoV-2 infection by a nucleic acid–positive test and were enrolled in this study. Patients were divided into two groups according to the clinical classification of patients with novel coronavirus infection, first group is mild/moderate ill and the second group is severe/critical ill. Patients were defined as mild ill if their clinical symptoms were mild, and there was no sign of pneumonia on imaging. Patients were defined as moderate ill if they get persistent high fever<3 days or (and) cough, shortness of breath, etc, but the respiratory rate (RR)<30 times /min, and the finger oxygen saturation when breathing air at rest>93%. The imaging features of COVID-19 infection pneumonia can be seen. Patients were defined as severely ill if they met the following criteria: (1) respiratory distress (respiratory rate ≥ 30 breaths/min); (2) pulse oxygensaturation ≤ 93% on room air; (3) low arterial oxygenation ratio (PaO2/fraction of inspired oxygen ≤ 300). Patients were defined as critically ill if the met the following criteria: (1) respiratory failure requiring a form of mechanical ventilation; (2) shock; (3) complications with other organ failure that require monitoring and treatment in the intensive care unit (ICU).

Data on basic information (age, gender, and comorbidities) and medical history of present illness (laboratory values, treatment, et al) on admission were collected from electronic medical records for each participant.

### 2. Sample collection, processing and isolation of immunocytes

Peripheral venous blood samples were collected from SARS-CoV-2 infected patients in Peking University Third Hospital immediately after confirmed with SARS-CoV-2 infection by a nucleic acid–positive test. Firstly, the samples were centrifuged at 2000g for 10 minutes. And then the plasma on the top layer of the EDTA Vacutainer tubes (BD, NJ, USA) was aliquoted and stored at −80°C. And the residual samples were processed for lysed red blood cells using RBC lysis Buffer (Biolegend, CA, USA). White blood cells were collected for immunophenotyping assay. For intracellular cytokine staining and ELISPOT assay, peripheral blood mononuclear cells (PBMCs) were obtained from white blood cells after density gradient centrifugation in Ficoll (Sigma, MO, USA)

### 3. Flow cytometry

Immune cells phenotyping (Panel S1), T cell senescence (Panel S2) and T cell activation (Panel S3) were determined using specific markers for innate leukocytes, T cells and cytokines. Briefly, 1×10^6^ white blood cells were stained with the Cellular Senescence Detection Kit (Dojindo Molecular Technologies, Gaithersburg, MD) according to the manufacturer’s instructions and cell surface molecules in Panel S1 and S2 were stained for 20 min in the dark at room temperature. SARS-CoV-2 specific or non-specific cytokines production by T cells was detected in Panel S3. For assessment of SARS-CoV-2 non-specific cytokines production, 1×10^6^ PBMCs were cultured with PMA (Biolegend, CA, USA) and Brefeldin A (Biolegend, CA, USA) for 5 hours. For assessment of SARS-CoV-2 specific cytokines production, 2×10^6^ PBMCs were cultured with 2 mg/mL SARS-CoV-2 S protein (MabTech, Stockholm, Sweden) for 24 hours, Brefeldin A (Biolegend, CA, USA) was added 5 h before cell collection. PBMCs in Panel S3 were stained with surface makers of T cells firstly. Afterwards, PBMCs were fixed and permeabilized using the Staining Buffer Kit (BioLegend CA, USA). Lastly, intracellular proteins were stained. Cytoflex cytometer (Beckman, CA, USA) and Kaluza analysis flow cytometry software v. 2.1.1. was used for all flow cytometric analyses. All antibodies in the three panels were listed in Supplementary Table 1. The gating strategy was shown in Supplementary Figure 1.

### 4. SARS-CoV-2 specific total IgG and IgM titer evaluation

96-well plate (Thermo Fisher, MA, USA) were coated with 1 µg/ml of WT spike protein, or WT and Omicron BF.7 RBD protein (Sino Biological, Beijing, China) at 4 °C overnight. Plates were then blocked with ELISA Assay Diluent (Biolegend, CA, USA) at room temperature for 2 h. Serum samples were serially diluted and added to the blocked plates before incubation at room temperature for an hour. After incubation, bound antibodies were either detected with goat anti-human IgM-HRP antibody (Sigma, SL, USA) for IgM assessment or goat anti-human IgG-HRP (Invitrogen, CA, USA) for IgG assessment. After 45 minutes, plates were developed by TMB substrate (Biolegend, CA, USA) and the reactions were stopped by adding ELISA stop solution (Solarbio, Beijing, China). The absorbance at 450 nm and 630 nm were measured with Spark reader (Tecan, Männedorf, Switzerland). The endpoint dilution titer was calculated in GraphPad Prism using a 0.15 OD 450-630 nm cutoff.

### 5. Neutralizing antibodies assay

SARS-CoV-2 RBD neutralizing antibody test kit (Vazyme, Nanjing, China) was used for determining RBD neutralizing antibodies inhibition rate. Initial antibody dilution rate was 1:10 by adding 8 μL sera into 72 μL dilution buffer. And the next assay was based on the protocol of manufacturer. Briefly, the ACE2 binding plate was incubated with plasma at 37 ℃ for 20 minutes. Then the plate was washed with wash buffer and 100 μL TMB was added followed by incubation for 15 minutes at 37 ℃. Then 50 μL stop solution was added and the absorbance at 450 nm were measured with Spark reader (Tecan, Männedorf, Switzerland). Inhibition (%) = (1 − sample OD450/negative control OD450) × 100%.

### 6. ELISPOT assay

Cellular specific immune responses in the patients were assessed using IFN-γ precoated ELISPOT kits (MabTech, Stockholm, Sweden), according to the manufacturer’s protocol. Briefly, the plates were blocked using RPMI 1640 (Hyclone, KCDC, USA) containing 10% FBS and incubated for 30 minutes. PBMCs were then plated at 3 × 10^5^ cells/well, stimulated with 2 mg/ml human peptide pool for SARS-CoV-2 S protein (MabTech, Stockholm, Sweden), PMA (Biolegend, CA, USA) was used as positive control and RPMI 1640 was used as negative control. After incubation at 37℃, 5% CO2 for 24 hours, plates were washed with wash buffer and biotinylated anti-human IFN-γ antibody was added followed by incubation for 2 hours at room temperature. Following the addition of AEC substrate solution, the numbers of spot-forming cells were counted using ELISPOT reader AID ELISPOT (AID, Strassberg, Germany).

### 7. Plasma soluble factors multiplex immune assay

53 plasma samples were analyzed by LEGENDPlex^TM^ using the Human Proinflammation Chemokine Panel 1, the Human Chemokine Panel 2 and Human CD8/NK Panel (Biolegend, CA, USA). The assay was performed according to the manufacturer’s instructions. Flow cytometric analysis was performed on CytoFLEX S (Beckman Coulter). Data were analyzed using online software (Biolegend).

### 8. Statistical analysis

GraphPad Prism 9 and SPSS 23 was used for graphic representation and statistical analysis. All reported probability values were two-tailed, and a P less than 0.05 was considered statistically significant. Statistical testing included t test (data conformed to the normal distribution), Man Whitney U test (data not conformed to the normal distribution), chi square (χ2) and Fisher’s exact tests, and Kaplan-Meier survival analysis with Gehan-Breslow-Wilcoxon test. Correlation were tested by Spearman’s rank coefficient (data not conformed to the normal distribution). Cut-off level (high vs. low) of Tsen was computed by log-rank maximization method.

### 9. Study Approval

All sampling and experimental steps in this study were approved by the Ethics Committee of Peking University Third Hospital (License No. IRB00006761-M2022865).

## Results

### 1. Patient clinical characteristics

Demographics characteristics and clinical features of patients were displayed in Table 1. The average age was 80.10 ± 9.89 years, and 64 (of 100; 64%) of them were men. The average BMI of patients was 23.81 ± 3.91 kg/m^2^; 38.5% (37 of 96) were overweight (24.0 ≤ BMI ≤ 27.9 kg/m^2^) and 12% (12.5 of 96) were obese (BMI ≥ 28.0 kg/m^2^) according to the Chinese BMI cutoffs. The most common comorbidities were hypertension (52 of 100; 52%), diabetes 25 of 100; 25%), and cardiovascular diseases (24 of 100;24%). cough (85 of 100; 85%), fever (82 of 100; 82%), sputum production (80 of 100; 80%) and dyspnea (60 of 100; 60%) were the most common symptoms at onset of illness. Compared with patients with mild/moderate illness, patients who were severe/critically ill were tend to more dyspneic (p=0.057). Other characteristics and symptoms had no significant difference between the 2 groups.

**Table 1.**
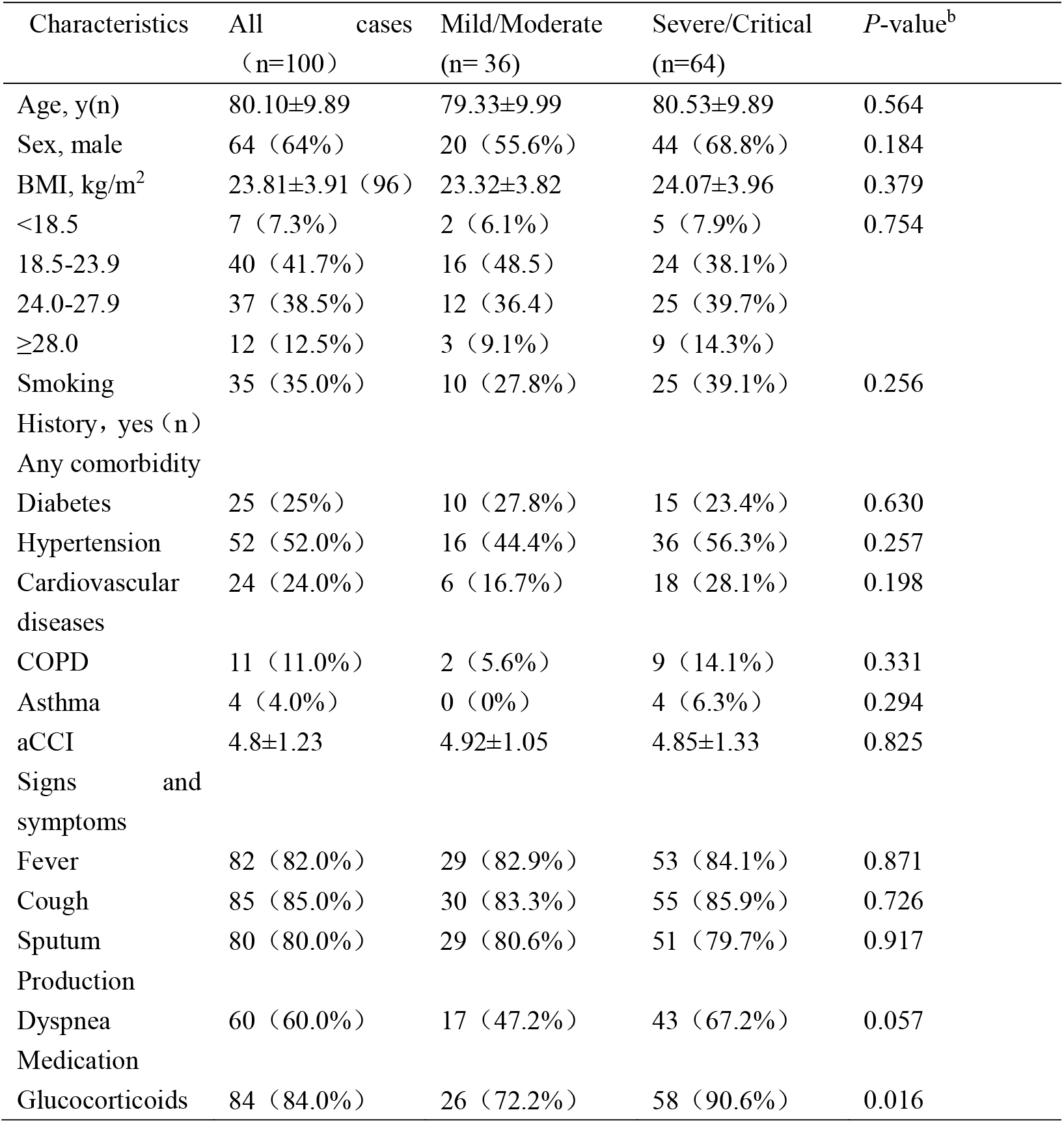
Demographics, Characteristics, and Clinical Features of Patients with Coronavirus Disease 2019^a^.

| Characteristics | All cases<br>(n=100) | Mild/Moderate<br>(n= 36) | Severe/Critical<br>(n=64) | P-value<sup>b</sup> |
| --- | --- | --- | --- | --- |
| Age, y(n) | 80.10±9.89 | 79.33±9.99 | 80.53±9.89 | 0.564 |
| Sex, male | 64 (64%) | 20 (55.6%) | 44 (68.8%) | 0.184 |
| BMI, kg/m<sup>2</sup> | 23.81±3.91 (96) | 23.32±3.82 | 24.07±3.96 | 0.379 |
| <18.5 | 7 (7.3%) | 2 (6.1%) | 5 (7.9%) | 0.754 |
| 18.5-23.9 | 40 (41.7%) | 16 (48.5) | 24 (38.1%) |  |
| 24.0-27.9 | 37 (38.5%) | 12 (36.4) | 25 (39.7%) |  |
| ≥28.0 | 12 (12.5%) | 3 (9.1%) | 9 (14.3%) |  |
| Smoking | 35 (35.0%) | 10 (27.8%) | 25 (39.1%) | 0.256 |
| History, yes (n) |  |  |  |  |
| Any comorbidity |  |  |  |  |
| Diabetes | 25 (25%) | 10 (27.8%) | 15 (23.4%) | 0.630 |
| Hypertension | 52 (52.0%) | 16 (44.4%) | 36 (56.3%) | 0.257 |
| Cardiovascular diseases | 24 (24.0%) | 6 (16.7%) | 18 (28.1%) | 0.198 |
| COPD | 11 (11.0%) | 2 (5.6%) | 9 (14.1%) | 0.331 |
| Asthma | 4 (4.0%) | 0 (0%) | 4 (6.3%) | 0.294 |
| aCCI | 4.8±1.23 | 4.92±1.05 | 4.85±1.33 | 0.825 |
| Signs and symptoms |  |  |  |  |
| Fever | 82 (82.0%) | 29 (82.9%) | 53 (84.1%) | 0.871 |
| Cough | 85 (85.0%) | 30 (83.3%) | 55 (85.9%) | 0.726 |
| Sputum Production | 80 (80.0%) | 29 (80.6%) | 51 (79.7%) | 0.917 |
| Dyspnea | 60 (60.0%) | 17 (47.2%) | 43 (67.2%) | 0.057 |
| Medication |  |  |  |  |
| Glucocorticoids | 84 (84.0%) | 26 (72.2%) | 58 (90.6%) | 0.016 |

BMI, body mass index; aCCI, age-adjusted Charlson Comorbidity Index.

Compared with patients with mild/moderate illness, more severe/critically ill patients were received glucocorticoids (84 of 100; 84%) during the entire hospital stay. The comparisons of treatment and medication between the 2 groups are shown in Table 1.

Laboratory characteristics of 100 patients were collected and are presented in Table 2. On admission, white blood cell counts were below the reference range in 2 (of 100; 2%) patients and above the reference range in 23 (of 100; 23%) patients. Neutrophil counts were higher in severely/critically ill patients than in mildly/moderately ill patients (p=0.018, respectively) and lymphocyte count were lower in severely/critically ill patients than in mild/moderate ill patients (p=0.004, respectively). The levels of coagulation function indexes such as D-dimer on admission were higher in severely/critically ill patients than in mildly/moderately ill patients (p=0.035, respectively). Regarding the inflammatory markers, procalcitonin (PCT) is higher in severely/critically ill patients than in mildly/moderately ill patients (p=0.046, respectively). No significant differences in C-reactive protein (CRP) levels, hemoglobin, serum albumin, total bilirubin, alanine aminotransferase, aspartate aminotransferase, creatinine and uric acid were observed between the 2 groups.

**Table 2.**
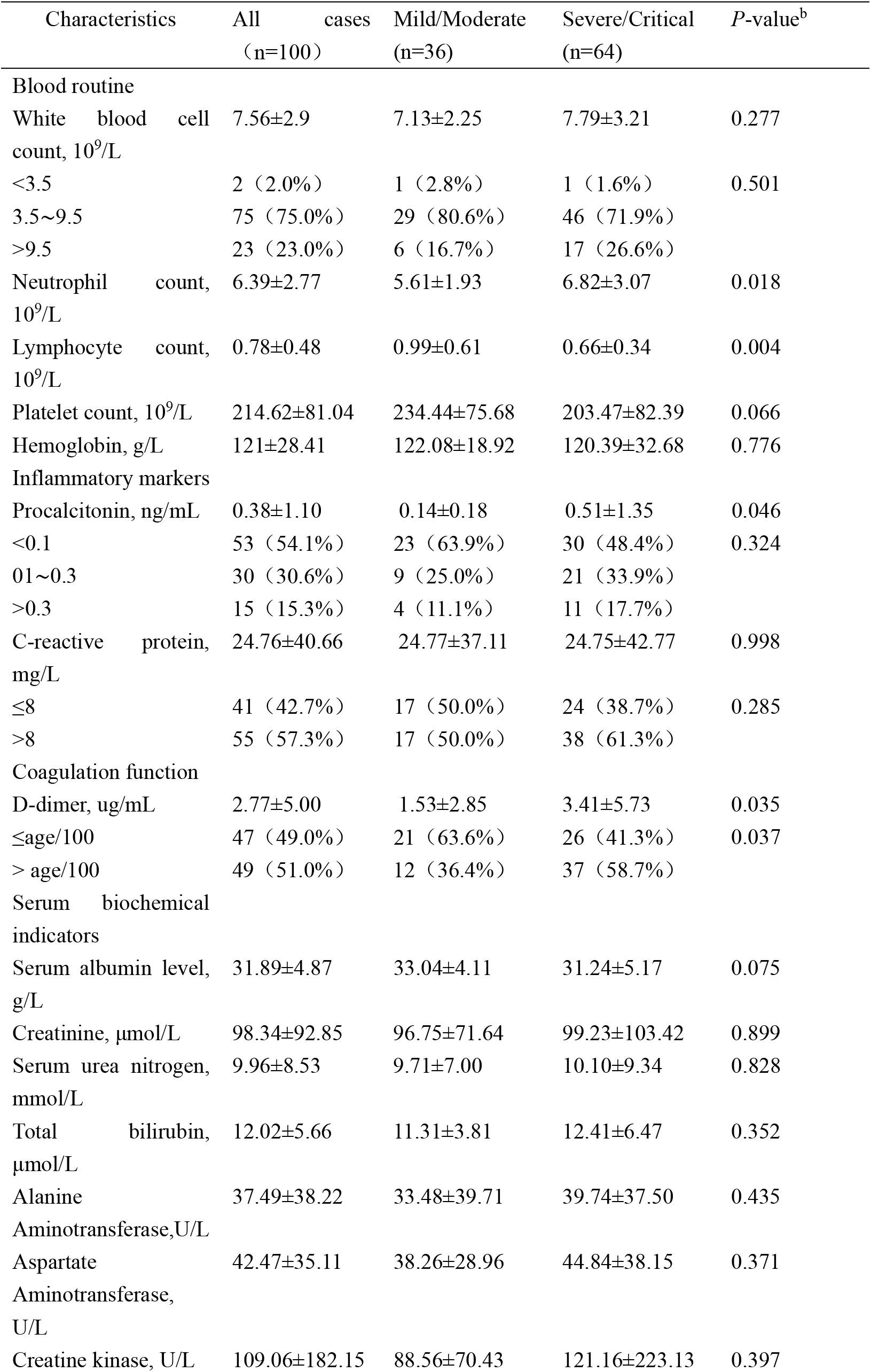
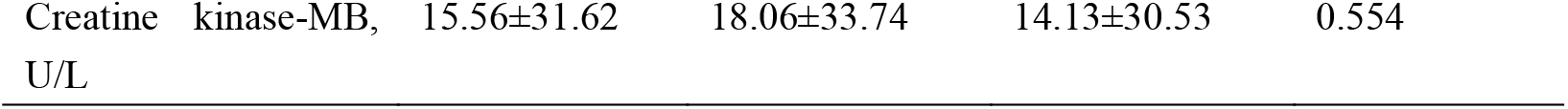
Laboratory Characteristics on Admission for Severely and Critically Ill Patients with Coronavirus Disease 2019^a^.

| Characteristics | All cases<br>(n=100) | Mild/Moderate<br>(n=36) | Severe/Critical<br>(n=64) | P-value<sup>b</sup> |
| --- | --- | --- | --- | --- |
| Blood routine |  |  |  |  |
| White blood cell count, 10<sup>9</sup>/L | 7.56±2.9 | 7.13±2.25 | 7.79±3.21 | 0.277 |
| <3.5 | 2 (2.0%) | 1 (2.8%) | 1 (1.6%) | 0.501 |
| 3.5~9.5 | 75 (75.0%) | 29 (80.6%) | 46 (71.9%) |  |
| >9.5 | 23 (23.0%) | 6 (16.7%) | 17 (26.6%) |  |
| Neutrophil count, 10<sup>9</sup>/L | 6.39±2.77 | 5.61±1.93 | 6.82±3.07 | 0.018 |
| Lymphocyte count, 10<sup>9</sup>/L | 0.78±0.48 | 0.99±0.61 | 0.66±0.34 | 0.004 |
| Platelet count, 10<sup>9</sup>/L | 214.62±81.04 | 234.44±75.68 | 203.47±82.39 | 0.066 |
| Hemoglobin, g/L | 121±28.41 | 122.08±18.92 | 120.39±32.68 | 0.776 |
| Inflammatory markers |  |  |  |  |
| Procalcitonin, ng/mL | 0.38±1.10 | 0.14±0.18 | 0.51±1.35 | 0.046 |
| <0.1 | 53 (54.1%) | 23 (63.9%) | 30 (48.4%) | 0.324 |
| 0.1~0.3 | 30 (30.6%) | 9 (25.0%) | 21 (33.9%) |  |
| >0.3 | 15 (15.3%) | 4 (11.1%) | 11 (17.7%) |  |
| C-reactive protein, mg/L | 24.76±40.66 | 24.77±37.11 | 24.75±42.77 | 0.998 |
| ≤8 | 41 (42.7%) | 17 (50.0%) | 24 (38.7%) | 0.285 |
| >8 | 55 (57.3%) | 17 (50.0%) | 38 (61.3%) |  |
| Coagulation function |  |  |  |  |
| D-dimer, ug/mL | 2.77±5.00 | 1.53±2.85 | 3.41±5.73 | 0.035 |
| ≤age/100 | 47 (49.0%) | 21 (63.6%) | 26 (41.3%) | 0.037 |
| > age/100 | 49 (51.0%) | 12 (36.4%) | 37 (58.7%) |  |
| Serum biochemical indicators |  |  |  |  |
| Serum albumin level, g/L | 31.89±4.87 | 33.04±4.11 | 31.24±5.17 | 0.075 |
| Creatinine, μmol/L | 98.34±92.85 | 96.75±71.64 | 99.23±103.42 | 0.899 |
| Serum urea nitrogen, mmol/L | 9.96±8.53 | 9.71±7.00 | 10.10±9.34 | 0.828 |
| Total bilirubin, μmol/L | 12.02±5.66 | 11.31±3.81 | 12.41±6.47 | 0.352 |
| Alanine Aminotransferase, U/L | 37.49±38.22 | 33.48±39.71 | 39.74±37.50 | 0.435 |
| Aspartate Aminotransferase, U/L | 42.47±35.11 | 38.26±28.96 | 44.84±38.15 | 0.371 |
| Creatine kinase, U/L | 109.06±182.15 | 88.56±70.43 | 121.16±223.13 | 0.397 |

### 2. Increased CD4^+^ Tsens is increased in patients with severe/critical COVID-19

Loss of CD28 and gain of CD57 are prominent markers of senescent T cells ^11^. Therefore, we used the markers CD28 and CD57 to identify four populations within T cells: CD28^+^CD57^-^ (Tn), CD28^-^CD57^-^ (Tdn), CD28^+^CD57^+^ (Tdp) and CD28^-^CD57^+^ (Tsen, gating strategy in Supplementary Figure 1). Several other senescence markers were also detected. Tsen had the highest SA-β-gal activity and the expression of KLRG-1 compared with Tn, Tdn and Tdp subsets (Supplementary Figure 2a, b). Phenotype analysis revealed that Tsen were predominantly EM (CCR7^-^CD45RA^-^) or EMRA (CCR7^-^CD45RA^+^), whereas Tn were more naïve (CCR7^+^CD45RA^+^) and CM (CCR7^+^CD45RA^-^) cells (Supplementary Figure 2c). In addition, the percentage of Tn was reversely associated with the percentage of Tsen and Tsen/Tn ratio in both CD4^+^ and CD8^+^ T cells (Supplementary Figure 2d). We also found a significant correlation between CD4^+^ Tsen and CD8^+^ Tsen (r=0.30, p=0.002, Supplementary Figure 2d), suggesting that the senescence of CD4^+^ and CD8^+^ T cells was synchronous.

To determine whether T cell senescence was associated with COVID-19 severity, we compared the frequency of senescent T cells in the mild/moderate group with that in the severe/critical group. No change in the percentage of CD8^+^ Tn, Tdn, Tdp and Tsen was noted in mild/moderate patients compared to severe/critical patients (Figure 1a). However, the percentage of CD4^+^ Tn was markedly reduced while the percentage of CD4^+^ Tsen was significantly increased, resulting in an increased Tsen/Tn ratio in severe/critical group compared to mild/moderate group (CD4^+^ Tn, p=0.0303; CD4^+^ Tsen, p=0.0401; CD4^+^ Tsen/Tn, p=0.0334; Figure 1a).

**Figure 1.**
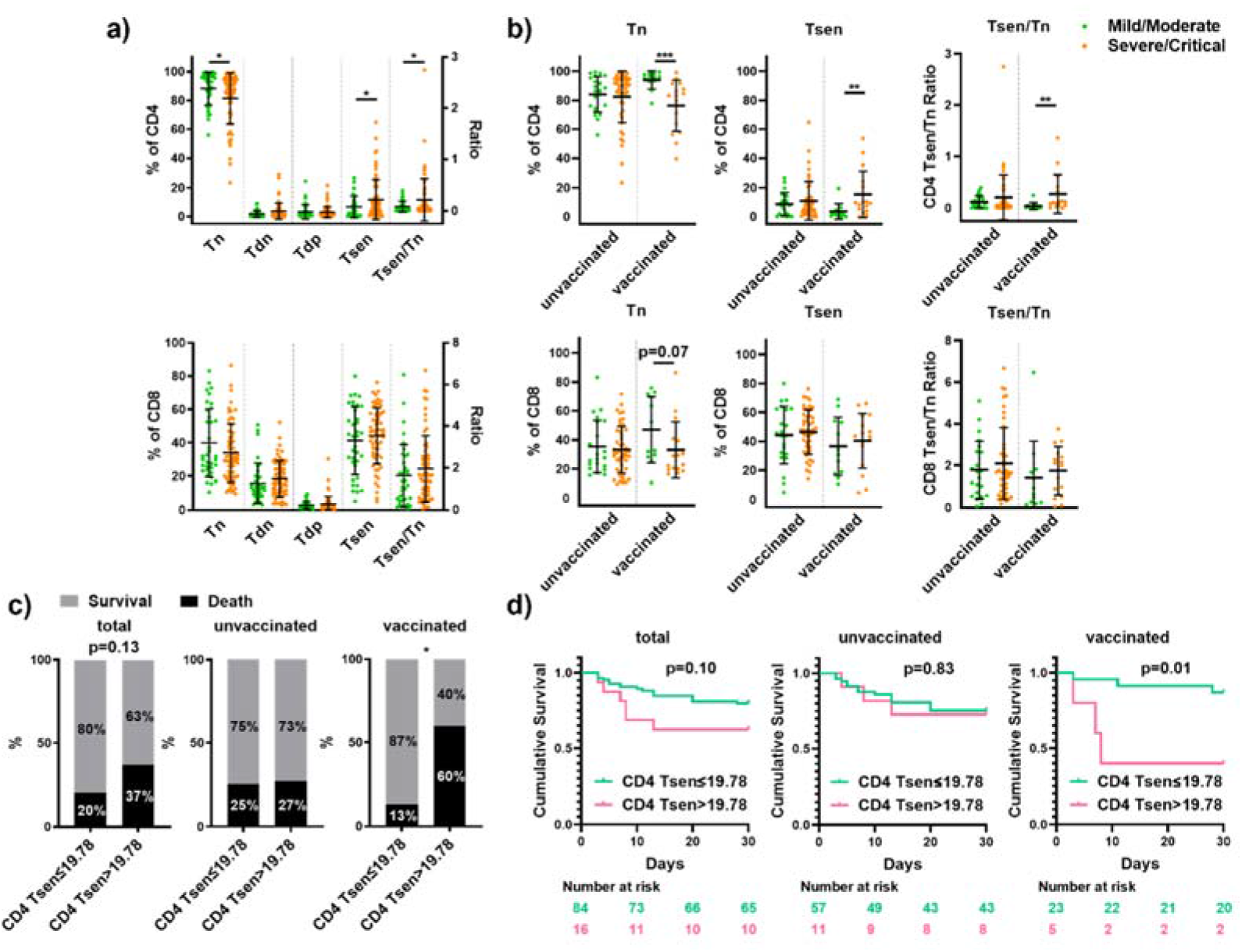
Increased CD4^+^ Tsens in severe/critical COVID-19. (a) The percentage of CD4^+^ and CD8^+^ T cell subsets: CD28^+^CD57^-^ (Tn), CD28^-^CD57^-^ (Tdn), CD28^+^CD57^+^ (Tdp), and CD28^-^CD57^+^ (Tsen) and Tsen/Tn ratio in mild/moderate patients (n=36) compared to sever/critical patients (n=64). (b) The percentage of CD4^+^ and CD8^+^ Tn, Tsen and Tsen/Tn ration in unvaccinated group (mild/moderate: n=22 versus sever/critical: n=46) and vaccinated group (mild/moderate: n=12 versus sever/critical: n=16). Groups were compared using Mann–Whitney U-test. Bars show mean with SD. (c) Survival rates in patients with COVID-19 stratified by levels of CD4 Tsen (the entire cohort: CD4 Tsen<19.78%, n=84 versus CD4 Tsen>19.78%, n=16; unvaccinated group: CD4 Tsen<19.78%, n=57 versus CD4 Tsen>19.78%, n=11; vaccinated group: CD4 Tsen<19.78%, n=23 versus CD4 Tsen>19.78%, n=5). P values for difference between survival rates were calculated using Fisher exact test. (d) Kaplan-Meier 30-day survival curves for COVID-19 patients with CD4 Tsen >19.78% versus CD4 Tsen <19.78% (the entire cohort: CD4 Tsen<19.78%, n=84 versus CD4 Tsen>19.78%, n=16; unvaccinated group: CD4 Tsen<19.78%, n=57 versus CD4 Tsen>19.78%, n=11; vaccinated group: CD4 Tsen<19.78%, n=23 versus CD4 Tsen>19.78%, n=5). P values for difference between two groups of curves were calculated by the log rank test. *, p< 0.05; **, p <0.01; ***, p <0.001.

To study the relationship between T cell senescence and vaccine efficacy, patients were further divided into unvaccinated (n=68) and vaccinated (n=28) groups. The association was confirmed in vaccinated group, where patients who had less Tn but more Tsen CD4^+^ T cells were more vulnerable to serious illness after a breakthrough infection (CD4^+^ Tn, p=0.0007; CD4^+^ Tsen, p=0.0044; CD4^+^ Tsen/Tn, p=0.0037, Figure 1b).

CD8^+^ Tn, Tsen and Tsen/Tn ratio also showed similar trend (CD8^+^ Tn, p=0.0661; CD8^+^ Tsen, p=0.6313; CD8^+^ Tsen/Tn, p=0.2053, Figure 1b), although not significant. Whereas, in unvaccinated patients, the percentage of Tn, Tsen as well as Tsen/Tn ratio were not associated with severity of COVID-19 (Figure 1b).

To further explore the correlation between T cells senescent with clinical characteristics, we examined a series of patients’ clinical indicators with circulating immune profiles. We found senescent T cells accumulated with age, obesity and age-related diseases, such as cardiovascular disease and chronic obstructive pulmonary disease (COPD). However, in the very old individuals (age > 80 yrs old), the percentage of Tn and Tsen was not affected by age, BMI and Charlson Comorbidity Index, which indicates that Tsen may reach the plateau stage (Supplementary Figure 3a). CD8^+^ Tsen was correlated positively with CRP (r=0.254, p=0.012) and LDH (lactate dehydrogenase; r=0.302, p=0.002) but inversely with PLT (platelet; r=-0.219, p=0.027). CD4^+^ Tn was positively correlated with Creatinine (r=0.225, p=0.029; Supplementary Figure 3a). To determine the relationship between systemic immune cell profile and T cell senescence, flow cytometric analysis was performed on circulating immune cells in COVID-19 patients. In accordance with previous report^12^, both the percentage and cell counts of lymphocytes, CD3^+^ T cells, CD4^+^ T cells, CD8^+^ T cells, B cells, NK cells, monocytes and DC (dendritic cells) showed statistically significant reduction in patients with severe/critical COVID-19 disease compared to mild/moderate disease (Supplementary Figure 3b,c). Moreover, CD8^+^ Tsen was inversely correlated with lymphocytes, T cells, B cells, monocytes and DC, whereas CD4^+^ Tsen was inversely correlated with monocytes and DC, which participated in antigen presenting (Supplementary Figure 3d,e). These findings suggested distinct contributions of CD4^+^ and CD8^+^ Tsen on the progression of severe disease.

Next, we evaluated the association of CD4^+^ Tsen with survival rate in COVID-19. The median value of CD4^+^ Tsen was 5.19% (min 0.01%, max 64.95%). A CD4^+^ Tsen cut-off (19.78%) was computed by log-rank maximization method (Supplementary Figure 4). 16% (16/100) patients had ≥ 19.78% CD28^-^CD57^+^ among CD4^+^ T cells. As compared to patients with low levels CD4^+^ Tsen (<19.78%), more patients in CD4^+^ Tsen high group had developed severe and critical illness (87% versus 60%, p=0.045). Other clinical characteristics of patients with high and low levels of CD4^+^ Tsen were comparable (Supplementary Table 2, 3). As shown in Figure 1, the survival rate was associated with peripheral blood levels of CD4^+^ Tsen, where patients with high levels (>19.78%) showed a decreased survival rate as compared to those with low levels (<19.78%, 80% vs. 63%, p=0.13), especially in the breakthrough infection (87% versus 40%, p=0.02, Figure 1c). However, the survival rate was comparable between two groups in unvaccinated patients (75% versus 73%, Figure 1c). The association was confirmed by the survival analysis using Kaplan-Meier estimate, which shows CD4^+^ Tsen may act as an efficient prognostic biomarker especially in the breakthrough infection (Figure 1d).

### 3. High senescent CD4^+^ T cells is correlated with lower spike-specific antibody level

Virus-neutralizing antibodies have been implied in protection against infection ^13, 14^. To test the relationship between T cell senescence and virus-specific antibody production, we measured IgG and IgM titers in the serum samples against the original SARS-CoV-2 strain (S1 and RBD, S receptor binding domain) and the Omicron variants BF.7 (RBD). Since the domestic epidemic variant at that time was BF.7, the anti-BF.7 RBD antibody titers were significantly higher compared to anti-WT S1 and anti-WT RBD antibody titers (supplementary figure 5a). Indeed, vaccinated patients had higher levels of anti-spike specific IgG and IgM, but no difference in anti-BF.7 RBD IgM (supplementary figure 5b,c), suggesting that vaccination provided limited protection targeting Omicron variants in older adults. In the entire cohort, the anti-WT S1 IgG titers in patients with CD4^+^ Tsen >19.78% (median, 89) were approximately 90% lower in median as compared with those in patients with CD4^+^ Tsen <19.78% (median, 870; Figure 2a). Moreover, the median IgG and IgM titers against BF.7 RBD were extremely dampened in CD4^+^ Tsen >19.78% group (Figure 2a, b). These findings were confirmed in both primary infections and breakthrough infections (Figure 2a,b).

**Figure 2.**
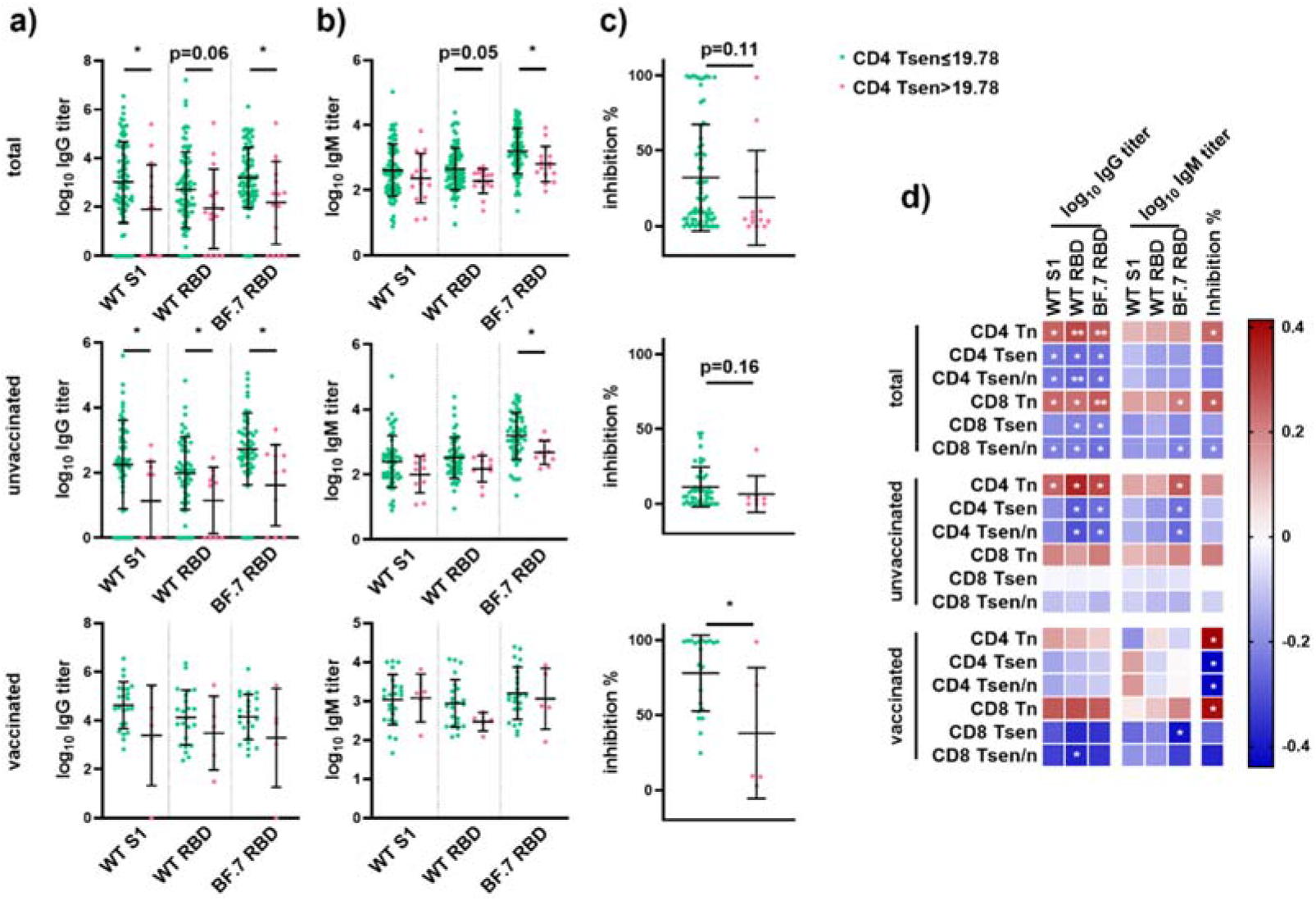
High senescent CD4^+^ T cells is correlated with lower spike-specific antibody level Plasma IgG. (a) and IgM (b) against the S1 domain of original SARS-CoV-2 strain (WT S1), the RBD domain of original strain (WT RBD) and the Omicron variants BF.7 (BF.7 RBD), and neutralization antibody inhibition rate (c) in COVID-19 patients with different level of CD4 Tsen (the entire cohort: CD4 Tsen<19.78%, n=80 versus CD4 Tsen>19.78%, n=15; unvaccinated group: CD4 Tsen<19.78%, n=53 versus CD4 Tsen>19.78%, n=10; vaccinated group: CD4 Tsen<19.78%, n=23 versus CD4 Tsen>19.78%, n=5). (d) Correlations between the percentage of senescent T cells and IgG and IgM titers (against WT S1, WT RBD and BF.7 RBD), inhibition rate (the entire cohort: n=95, unvaccinated group: n=63; vaccinated group: n=28). Statistical comparisons across cohorts were performed using the Mann-Whitney test. Spearman’s rank correlation was used to identify relationships between two variables. R values are indicated by color. Significant correlations were indicated by white asterisks. *, p< 0.05; **, p <0.01.

We next went on to assess the functional activity of spike-specific antibodies through measurement of neutralizing activity against ancestral SARS-CoV-2 variants. The inhibition rate was higher in vaccinated patients than that in unvaccinated ones (70.28%±32.74% versus 10.68%±13.08%, p<0.0001, Supplementary Figure 5d), indicating that neutralizing activity is relatively enhanced following vaccination. However, in vaccinated subgroup, neutralization of ancestral virus was markedly impaired in patients with CD4^+^ Tsen >19.78% compared with patients with CD4^+^ Tsen <19.78% (37.99%±43.59% versus 77.97%±25.25%, p=0.049, Figure 2c). The same trend was also found in the entire cohort as well as the unvaccinated subgroup (Figure 2c).

We also examined the association of spike-specific antibodies and T cell senescence. The same trend was found in the whole population, the unvaccinated group and the vaccinated group. In the whole cohort, anti-spike-specific IgG, anti-RBD IgG and anti-BF.7 RBD IgG levels were negatively correlated with senescent CD4^+^ and CD8^+^ T cells (Figure 2d). Importantly, in unvaccinated patients, anti-BF.7 RBD IgG and IgM levels were strongly reversely correlated with CD4^+^ Tsen. Moreover, neutralizing activity showed a positive relationship with CD4^+^ Tn, but a negative relationship with CD4^+^ Tsen and this phenomenon was more obvious in vaccinated group (Figure 2d).

Overall, these results above indicated that the accumulation of senescent CD4^+^ T cells may impair the production and neutralizing activity of spike-specific antibodies, which may further accelerate the severity and mortality in COVID-19 older patients.

### 4. Higher granzyme B and lower IL-2 is associated with defect in antibody production

To further characterize the phenotypic function of CD4^+^ Tsen, we made the comparison of the cytokine production between different amount of CD4^+^ Tsen groups. The data revealed that patients in CD4^+^ Tsen high group had less IL-2^+^CD4^+^ T cells but more granzyme B^+^ CD4^+^T cells compared to patients in CD4^+^ Tsen low group (Figure 3a). In line with these findings, we also observed a remarkable elevation of granzyme B^+^ T cells but a reduction of IL-2^+^ T cells in patients with severe/critical COVID-19 compared to patients with mild/moderate illness (supplementary figure 6a). In addition, the amount of both CD4^+^ and CD8^+^ Tsen were positively correlated with the percentage of granzyme B producing T cells, and negatively correlated with the frequency of IL-2 producing T cells (supplementary figure 6b). Cytokine production profiles analysis showed that compare to Tn, Tdn and Tdp, Tsen preferred to produce granzyme B, TNF-α and IFN-γ, but the production of IL-2 was impaired (supplementary figure 6c). Significantly, the percentage of IL-2^+^ T cells was positively correlated with the production and neutralizing activity of spike-specific antibodies (Figure 3b), indicating that the defect in IL-2 production of CD4^+^ Tsen may be responsible for their lower IgG responses.

**Figure 3.**
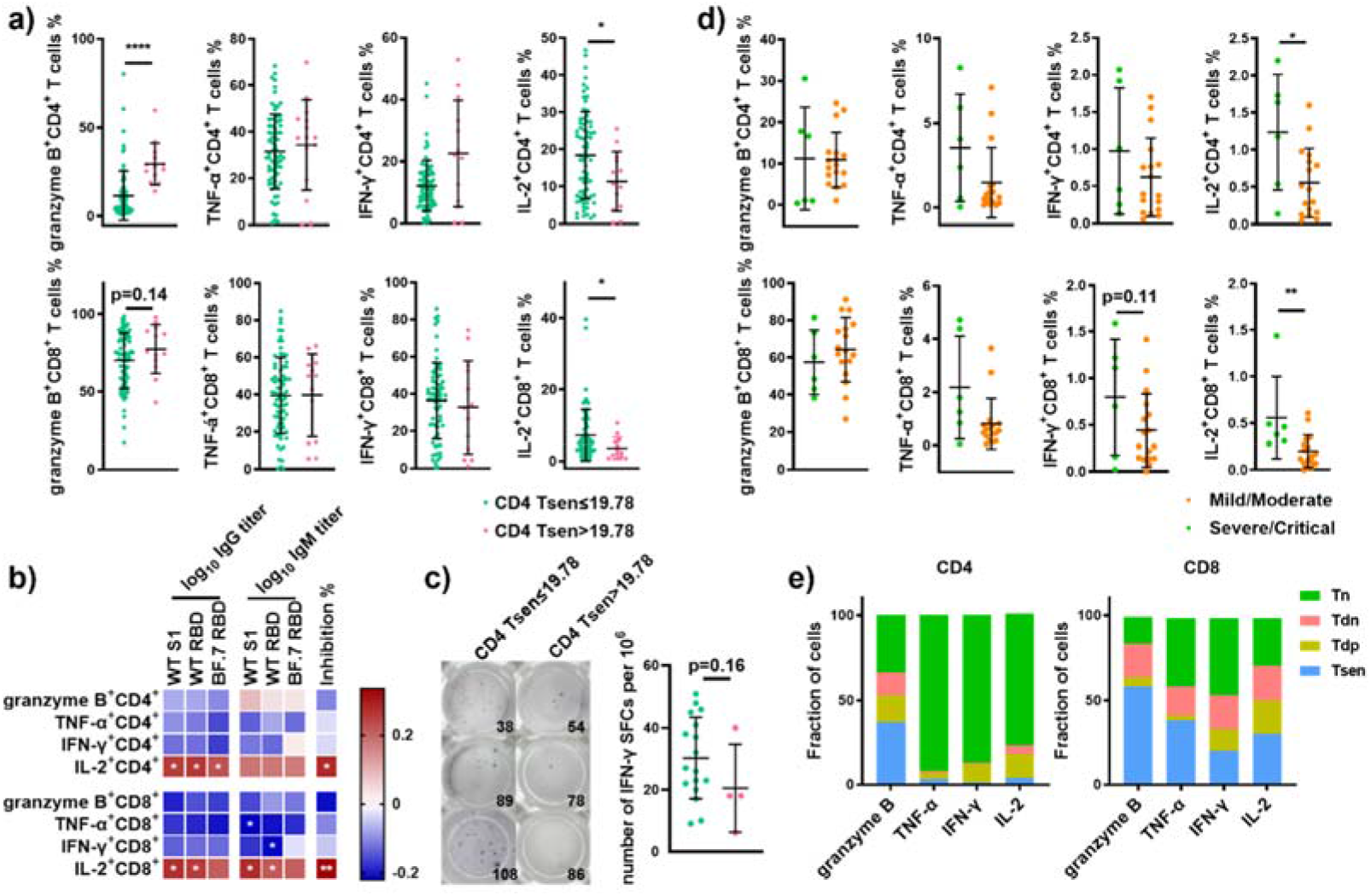
Higher granzyme B^+^ and lower IL-2^+^ T cells is associated with defect in antibody level. (a) Frequency of cytokine^+^ T cells (after PMA stimulation) in patients with CD4 Tsen <19.78% group (n=74) and CD4 Tsen >19.78% group (n=13). (b) Correlations between the percentage of cytokine^+^ T cells and plasma IgG and IgM against the S1 domain of original SARS-CoV-2 strain (WT S1), the RBD domain of original strain (WT RBD) and the Omicron variants BF.7 (BF.7 RBD, n=83), inhibition rate (n=74). (c) Frequency of IFN-γ SFCs after stimulation with Spike in patients with CD4 Tsen <19.78% group (n=17) and CD4 Tsen >19.78% group (n=4). The number represents the patient number. (d) Frequency of cytokine^+^ T cells (after Spike stimulation) in patients with mild/moderate illness (n=6) and severe/critical illness (n=17). (e) The frequency of Tn, Tdn, Tdp and Tsen (defined using the markers CD57 and CD28) within spike-specific cytokine^+^ T cells (n = 21). Groups were compared using Mann–Whitney U-test. Bars show mean with SD. Spearman’s rank correlation was used to identify relationships between two variables. R values are indicated by color. Significant correlations were indicated by white asterisks. *, p< 0.05; **, p <0.01; ****, p <0.0001.

Then, we measured SARS-CoV-2-specific T cell responses by stimulating PBMCs with pools of overlapping peptides spanning the SARS-CoV-2 S1 and S2 subunits of the Spike protein using interferon-γ (IFN-γ) enzyme-linked immunospot (ELISpot) assay. The overall Spike-specific T cell response in patients with CD4^+^ Tsen >19.78% was slightly lower than in patients with CD4^+^ Tsen <19.78% (18 versus 27, p=0.16; Figure 3c). Since the ELISpot assay does not allow identification of T cell subsets, we utilized intracellular cytokine staining by flow cytometry to further characterize the phenotype of responding cells. PBMCs were stimulated with the Spike peptide pool and CD4^+^ and CD8^+^ T cells were analyzed for the production of granzyme B, tumor necrosis factor-α (TNF-α), IFN-γ and interleukin-2 (IL-2). The gating strategy is depicted in supplementary Figure 1. Due to the small numbers, we could not compare CD4^+^ Tsen>19.78% (n=2) to CD4^+^ Tsen<19.78% (n=21). Instead, we investigated whether there was a difference between mild/moderate group and severe/critical group. In agreement with the results obtained with the IFN-γ ELISpot assay, IFN-γ and TNF-α-producing CD4^+^ and CD8^+^ T cells showed a decreased tendency in severe/critical illness compared to mild/moderate illness (Figure 3d). Similarly, the production of IL-2 in spike-specific CD4^+^ and CD8^+^ T cells was significantly higher in mild/moderate illness (Figure 3d). No differences were found between severity of illness in relation to T cells producing granzyme B (Figure 3d). Phenotype analysis revealed that the majority of IL-2^+^, TNF-α^+^ and IFN-γ^+^ spike specific T cells were Tn, whereas most of granzyme B^+^ spike-specific T cells were Tsen (Figure 3e).

Taken together, our findings suggested that higher granzyme B and lower IL-2 is the distinct feature of CD4^+^ Tsen, which is associated with defect in spike-specific antibody production.

### 5. Tsens and spike-specific antibodies is associated with plasma inflammatory factors

Finally, plasma level of 39 soluble factors were detected by CBA in 53 COVID-19 patients. Consistent with previous report, severe/critical patients showed increased plasma level of IL-6 compared to mild/moderate patients (71.54±23.36 pg/ml versus 22.02±9.565 pg/ml, p=0.130, Supplementary Table 4). No other significant differences were found in our cohort, indication that as the key component of SASP (Senescent associated secretory phenotype), IL-6 may mediate important inflammatory role in COVID-19 progression.

Next, we analyzed potential correlations among spike-specific antibody levels and plasma soluble factors. Plasma level of cytokines and chemokines have been grouped according to one of their main functions, as reported in (Figure 4a). As expected, IL-2 was positively related to spike-specific antibody. Similarly, T cell related cytokines, IL-4 and IL-17A, as well as pro-inflammatory cytokines, IL-1α, IL-1β and IL-11, were positively correlated with levels of spike-specific IgG and IgM. These cytokines were involved in regulating T cell activation, inducing Th2 polarization and stimulating B cell antibody production^15–17^. However, we observed spike-specific antibodies were reversely correlated with IL-18 and IL-22. Regarding chemokines, spike specific antibodies, especially anti-BF.7 RBD IgM, was significantly associated with CXCL5, but negative associated with CCL4, CXCL9, CXCL10, CXCL11, which were reported to increase with disease severity ^18^. In addition, anti-viral factors exhibited different relationship with spike-specific antibodies. Levels of spike-specific antibody was positively correlated with IFN-γ and granzyme A and negatively correlated with granulysin and perforin, indicating that cellular immunity and humoral immunity may function in different ways.

**Figure 4.**
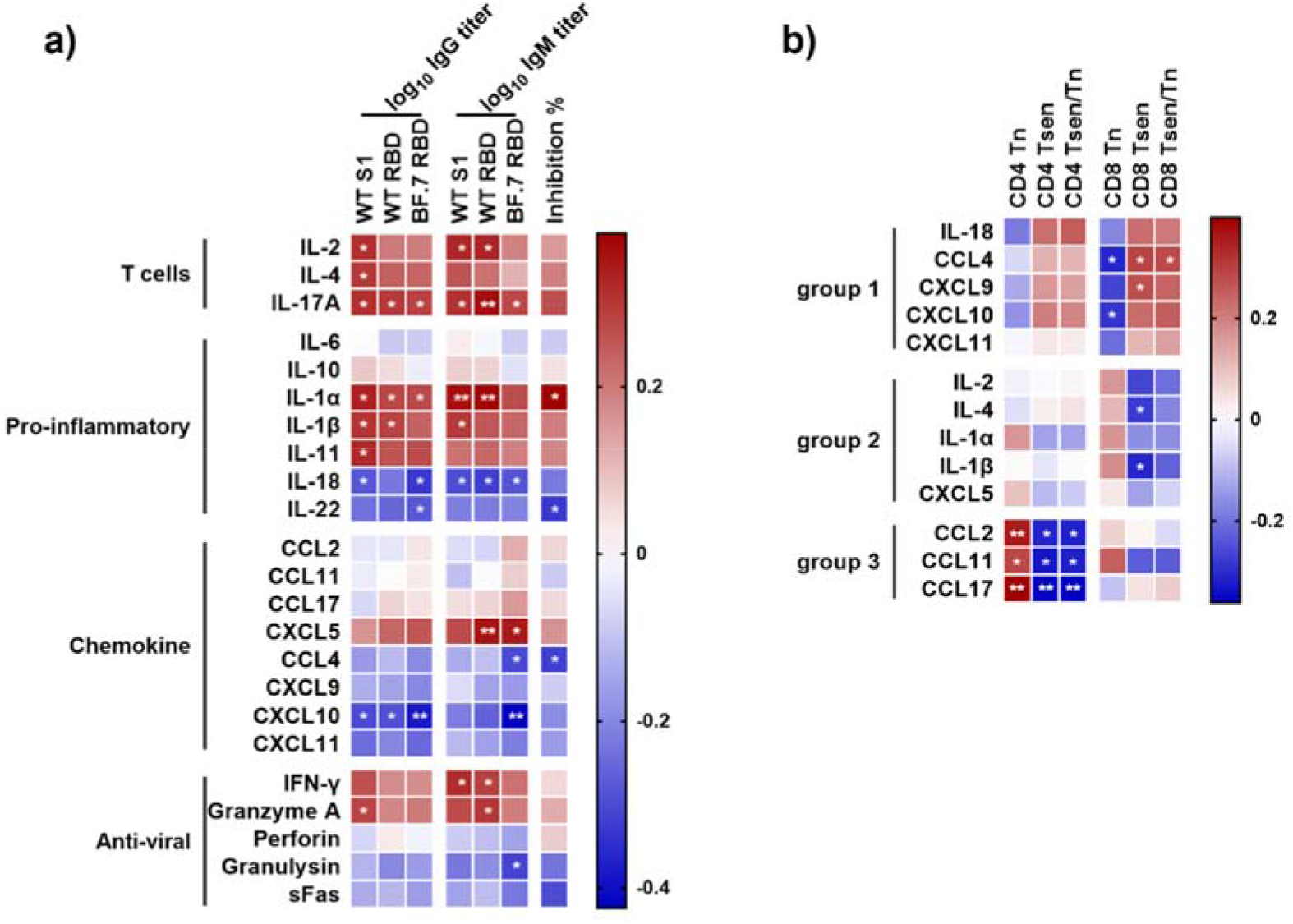
The correlation between plasma level of soluble factors and spike-specific antibody levels, neutralization ability as well as the percentage senescent T cells. (a) Correlation analysis between IgG and IgM titers (against WT S1, WT RBD and BF.7 RBD), inhibition rate and plasma levels of soluble factors (IL-2, IL-4, IL-17A, IL-6, IL-10, IL-1α, IL-1β, IL-11, IL-18, IL-22, CCL2, CCL11, CCL17, CXCL5, CCL4, CXCL9, CXCL10, CXCL11, IFN-γ, granzyme A, perforin, granulysin, sFas). The function mediated by each plasmatic molecule was indicated on the left. (b) Correlations between the percentage of senescent T cells and plasma levels of soluble factors (group 1: IL-18, CCL4, CXCL9, CXCL10, CXCL11; group 2: IL-2, IL-4, IL-1α, IL-1β, CXCL5; group 3: CCL2, CCL11, CCL17). Data were collected from 53 COVID-19 infected patients, except the inhibition rate was detected in 42 patients. Spearman’s rank correlation was used to identify relationships between two variables. R values are indicated by color. Significant correlations were indicated by white asterisks. *, p<0.05; **, p<0.01.

Moreover, we examined relationships between levels of plasma soluble molecules and T cell senescence. As shown in (Figure 4b), several paired parameters were identified with 3 types of significant correlations. Molecules in group 1, such as IL-18, CCL4, CXCL9, CXCL10 and CXCL11, showed negative relations with spike-specific antibody levels, but had positive relations with the frequency of Tsen in the peripheral blood. Molecules in group 2, such as IL-2, IL-4, IL-1α, IL-1β and CXCL5, were positively correlated with spike-specific antibody levels, while negatively correlated with the percentage of Tsen. Molecules in group 3, including CCL2, CCL11 and CCL17, exhibit a remarkable positive correlation with CD4^+^ Tsen, whereas had no significant connection with spike-specific antibody production. These results indicated that molecules in group 1 and group 2 may be involved in the decreased ability of senescent T cells in helping antibody production, while group 3 molecules may increase the risk of severe disease in other ways.

## Discussion

In this study, we investigated the role of T cell senescence in elderly patients with primary and breakthrough COVID-19 infections. We demonstrated that elderly patients with severe/critical illness had higher CD4^+^ Tsen compared with elderly patients with mild/severe disease. And the mortality rate was higher in patients with CD4^+^ Tsen > 19.78%. This phenomenon was more remarkable in vaccinated individuals. The percentage of CD4^+^ Tsens was correlated inversely with spike-specific IgG and IgM levels and neutralization ability. Moreover, IL-2 producing T cells and plasma levels of IL-2 was positively correlated with antibody levels. Our data for the first time illustrated that the percentage of CD4^+^ Tsen in the peripheral blood may act as an efficient biomarker for predicting the severity and prognosis of COVID-19 in older patients, especially in the breakthrough infection.

Immune responses to SARS-CoV-2 or vaccination are impacted by aging, CMV infection, as well as age related disease such as obesity, cardiometabolic diseases and frailty^7, 19, 20^. However, in our cohort, these clinical parameters were not correlated with disease severity or vaccine efficacy (Table 1), suggesting that other factors may be responsible for the individual variability in the outcome of COVID-19. We found an accumulation of CD4^+^ Tsen in severe/critical patients compared to mild/moderate patients (Figure 1a,b) and the death rate was significantly elevated in elderly patients with CD4^+^ Tsen > 19.78% (Figure 1c). Importantly, aging, cytomegalovirus (CMV) infection, as well as age related disease are reported to accelerate the senescence of T cells^7, 21^. Moreover, shorter leukocyte telomere length, a hallmark of biological aging, was associated with worse COVID-19 outcomes^1^. Our results indicated that for individuals in the later years of life, the senescence of CD4^+^ T cells might be one of the reasons why age or age-related disease is more likely to cause sever COVID-19.

The accumulation of senescent T cells is more pronounced for CD8^+^ T cells than CD4^+^ T cells with advanced age and CMV infection^22 23^. Therefore, when compared to young individuals, senescent SARS-CoV-2-Reactive CD8^+^ T Cells accumulated in elderly individuals, whereas CD4^+^ T cells proliferation and Th1 cytokine production upon COVID-19 recognition was comparable between elderly and young individuals ^19^. However, we found an accumulation of CD4^+^ Tsen, but not CD8^+^ Tsen, in the peripheral blood of severe/critical patients and was positively correlated with death rate (Figure 1a). These data suggested that within elderly patients, the defect in CD4^+^ Tsen may be responsible for immune responsiveness to COVID-19 infection and vaccination.

Both cellular and humoral immunity are involved in antiviral response against COVID-19^24, 25^. Humoral immunity is mediated by antibodies and memory B cells. Antibodies block infection by binding virus and preventing viral entry into host cells. Cellular immunity includes helper CD4^+^ T cells and cytotoxic CD8^+^ T cells. Since T cells do not recognize viruses until they have entered the host cell, T cells cannot prevent host cells from initially becoming infected, but they can respond rapidly once infection has occurred to limit virus replication and spread. Emerging evidence supports that immune responses that operate rapidly and efficiently after initial infection could prevent progression to severe disease^26^. The spike T cell responses peaked and remained unchanged after the first time of spike exposures ^27^, whereas the peak levels of anti-RBD antibody were observed after the second dose of inactivated, mRNA or adenovirus vaccination^10, 27–29^. These studies indicate the critical role of T cell immunity for long-term protection by COVID-19 vaccines. In our cohort, the spike-specific antibody titers were higher in vaccinated group than unvaccinated group (supplementary figure 5b-d). The quantity and quality of spike-specific antibody as well as the frequency of IL-2^+^ spike-specific CD4^+^ and CD8^+^ T cells was negatively correlated with CD4^+^ Tsen (Figure 2d, 3d). These findings suggest the COVID-19 vaccines could prevent progression to severe disease in elderly individuals with low percentage of CD4^+^ Tsens. In agreement, we found that the predictive effect of CD4^+^ Tsen on severe/critical disease was more significant in the vaccine group (Figure 1). Our findings also indicate that for elderly individuals with high levels of CD4^+^ Tsen, early antiviral therapy or the direct neutralizing antibody administration might be the alternative therapeutics approach.

There are several reasons for the susceptible to severe COVID-19 in elderly patients with accumulated Tsen. On one hand, insufficient epitope specific T cell clones may be responsible for impaired cellular immunity to COVID-19 infection in old adults. TCRβ diversity roughly declines linearly with age^30^ and the efficiency of TCR signaling is compromised in senescent T cells^31^. Compared to younger peoples (median age 63), the spike-specific cellular immune responses in older donors (median age 83) were impaired following vaccination^8^. Less CD8^+^ T cell clone expansion specific to SARS-CoV-2^10^ as well as lower induction and early contraction of CD4^+^ T cell responses were also reported in older adults^29^. In accordance with these reports, a decline in spike-specific IFN-γ and IL-2 producing T cells was found in patients with elevated CD4^+^ Tsen (Figure 3c,d). Although Tsen are more capable of producing granzyme B than other subsets (Figure 3a, supplementary figure 6b,c), our data suggest that granzyme B alone may serve as two-edged sword, killing the virus, but aggravating pulmonary damage and fibrosis^32^.

On the other hand, T cell senescence may affect the production of COVID-19 specific antibodies. Recent studies showed that vaccine-induced Spike-specific antibody was inversely correlated with senescent CD8^+^ T cells^7,9^. Likewise, we found that spike-specific antibody titer and neutralization ability was negatively associated with CD4^+^ Tsen in old adults (Figure 2), indicating that CD4^+^ Tsen may have defect in promoting B cell antibody production. We and others reported a decline of IL-2 production in aged patients ^9, 33^ or patients with CD4^+^ Tsen > 19.78% (Figure 3a). IL-2 enhances T cell activation by stimulating expansion and differentiation of conventional T cells^34^, which may delay the onset of lymphopenia for COVID-19 patients^35, 36^. T cells derived IL-2 was also able to promote the proliferation and antibodies production of B cells ^37^. Currently, one clinical trial is investigating the potential of administrating of IL-2 in the treatment of COVID-19^38^.

Besides the findings described above, our data has to be interpreted carefully due to some limitations. First, our clinical studies were conducted in a single institution and the sample size was limited. Therefore, the cut-off value (19.78%) can be varied due to the population size. Second, we did not detect the senescence and function of antigen-presenting cells and B cells that are also critical for vaccine-induced immunity, although it is reported that they are less affected by aging than T cells^39^. Third, we provided evidence only of associations between CD4^+^ Tsen responses with antibody and COVID-19 infection severity. Further studies are warranted to investigate causal relationships among these parameters.

## Conclusions

In conclusion, we demonstrated the accumulation of CD4^+^ Tsen, with defect in IL-2 production, was negatively correlated with the quantity and quality spike-specific antibody production, potentially enabling progression to severe outcome. Our study highlights CD4^+^ Tsen as a better indicator than other clinical parameters for immune response to COVID-19 infection in older adults. These findings suggest that vaccines which provoke CD4^+^ specific immunity might be efficacious for individuals in the later years of life or the direct neutralizing antibody administration for the high CD4^+^ Tsen patients might be the alternative therapeutics approach.

## Abbreviations

ACE: angiotensin converting enzyme

BMI: body mass index

CD: cluster of differentiation

COVID: corona virus disease

CBA: cytometric bead array

CRP: C-reactive protein

CM: central memory

COPD: chronic obstructive pulmonary disease

CXCL: C-X-C motif ligand

CCL: C-C motif ligand

DC: dendritic cells

ELISA: enzyme-linked immunosorbent assay

Elispot: enzyme-linked immunoblot

EM: effector memory

EMRA: terminally differentiated effector cells

EDTA: ethylene diamine tetraacetic acid

IL: interleukin

ICU: intensive care unit

IgG: immunoglobulin G

IgM: immunoglobulin M

IFN-γ: interferon-γ

LDH: lactate dehydrogenase

PCT: procalcitonin

PLT: platelet

PBMCs: peripheral blood mononuclear cells

RBD: receptor binding domain

RBC: red blood cell

RR: respiratory rate

SARS-CoV-2: severe acute respiratory syndrome coronavirus 2

SASP: senescent associated secretory phenotype

Tsens: senescent T cells

TNF-α: tumor necrosis factor

Th: helper T cells

WT: wild type

## Author contributions

LXX, YCS, JZ and CC contributed to the concept development and study design. JZ, ZYL, ZNY, TZ, XXY and XXD performed the laboratory studies. TTH, YL, YLD, XYG, XL, JQR, YFR, JW, CC and YCS collected the clinical data. JZ, ZYL, TTH, XLL, JLY and LXX contributed to data analysis, figure preparation and drafted the manuscript. All authors read and approved the final manuscript.

## Acknowledgements

The authors acknowledge the contribution of all investigators at the participating study sites.

## References

1 Zhang, J. et al. Changes in contact patterns shape the dynamics of the COVID-19 outbreak in China. Science 368, 1481–1486, doi:10.1126/science.abb8001(2020).

2 Shahid, Z. et al. COVID-19 and Older Adults: What We Know. J Am Geriatr Soc 68, 926–929, doi:10.1111/jgs.16472 (2020).

3 Ramasamy, M. N. et al. Safety and immunogenicity of ChAdOx1 nCoV-19 vaccine administered in a prime-boost regimen in young and old adults (COV002): a single-blind, randomised, controlled, phase 2/3 trial. Lancet 396, 1979–1993, doi:10.1016/S0140-6736(20)32466-1 (2021).

4 Anderson, E. J. et al. Safety and Immunogenicity of SARS-CoV-2 mRNA-1273 Vaccine in Older Adults. N Engl J Med 383, 2427–2438, doi:10.1056/NEJMoa2028436 (2020).

5 Oyebanji, O. A., Mylonakis, E. & Canaday, D. H. Vaccines for the Prevention of Coronavirus Disease 2019 in Older Adults. Infect Dis Clin North Am 37, 27–45, doi:10.1016/j.idc.2022.11.002 (2023).

6 De Biasi, S. et al. Marked T cell activation, senescence, exhaustion and skewing towards TH17 in patients with COVID-19 pneumonia. Nat Commun 11, 3434, doi:10.1038/s41467-020-17292-4 (2020).

7 Semelka, C. T. et al. Frailty impacts immune responses to Moderna COVID-19 mRNA vaccine in older adults. Immun Ageing 20, 4, doi:10.1186/s12979-023-00327-x (2023).

8 Parry, H. et al. Vaccine subtype and dose interval determine immunogenicity of primary series COVID-19 vaccines in older people. Cell Rep Med 3, 100739, doi:10.1016/j.xcrm.2022.100739 (2022).

9 Palacios-Pedrero, M. A. et al. Signs of immunosenescence correlate with poor outcome of mRNA COVID-19 vaccination in older adults. Nature Aging 2, 896-+, doi:10.1038/s43587-022-00292-y (2022).

10 Xiao, C. C. et al. Insufficient epitope-specific T cell clones are responsible for impaired cellular immunity to inactivated SARS-CoV-2 vaccine in older adults. Nature Aging, doi:10.1038/s43587-023-00379-0 (2023).

11 Lee, Y. H. et al. Senescent T Cells Predict the Development of Hyperglycemia in Humans. Diabetes 68, 156–162, doi:10.2337/db17-1218 (2019).

12 Huang, W. et al. Lymphocyte Subset Counts in COVID-19 Patients: A Meta-Analysis. Cytometry A 97, 772–776, doi:10.1002/cyto.a.24172 (2020).

13 Jansen, J. M., Gerlach, T., Elbahesh, H., Rimmelzwaan, G. F. & Saletti, G. Influenza virus-specific CD4+ and CD8+ T cell-mediated immunity induced by infection and vaccination. J Clin Virol 119, 44–52, doi:10.1016/j.jcv.2019.08.009 (2019).

14 Kreye, J. et al. A Therapeutic Non-self-reactive SARS-CoV-2 Antibody Protects from Lung Pathology in a COVID-19 Hamster Model. Cell 183, 1058–1069 e1019, doi:10.1016/j.cell.2020.09.049 (2020).

15 Nakae, S., Asano, M., Horai, R. & Iwakura, Y. Interleukin-1 beta, but not interleukin-1 alpha, is required for T-cell-dependent antibody production. Immunology 104, 402–409, doi:10.1046/j.1365-2567.2001.01337.x (2001).

16 Nakae, S., Asano, M., Horai, R., Sakaguchi, N. & Iwakura, Y. IL-1 enhances T cell-dependent antibody production through induction of CD40 ligand and OX40 on T cells. J Immunol 167, 90–97, doi:10.4049/jimmunol.167.1.90 (2001).

17 Fung, K. Y. et al. Emerging roles for IL-11 in inflammatory diseases. Cytokine 149, 155750, doi:10.1016/j.cyto.2021.155750 (2022).

18 Liao, M. et al. Single-cell landscape of bronchoalveolar immune cells in patients with COVID-19. Nat Med 26, 842–844, doi:10.1038/s41591-020-0901-9 (2020).

19 Jo, N. et al. Aging and CMV Infection Affect Pre-existing SARS-CoV-2-Reactive CD8(+) T Cells in Unexposed Individuals. Front Aging 2, 719342, doi:10.3389/fragi.2021.719342 (2021).

20 Retuerto, M. et al. Shorter telomere length is associated with COVID-19 hospitalization and with persistence of radiographic lung abnormalities. Immun Ageing 19, 38, doi:10.1186/s12979-022-00294-9 (2022).

21 Zhang, J., He, T., Xue, L. & Guo, H. Senescent T cells: a potential biomarker and target for cancer therapy. EBioMedicine 68, 103409, doi:10.1016/j.ebiom.2021.103409 (2021).

22 Martinez-Zamudio, R. I. et al. Senescence-associated beta-galactosidase reveals the abundance of senescent CD8+ T cells in aging humans. Aging Cell 20, e13344, doi:10.1111/acel.13344 (2021).

23 Wertheimer, A. M. et al. Aging and cytomegalovirus infection differentially and jointly affect distinct circulating T cell subsets in humans. J Immunol 192, 2143–2155, doi:10.4049/jimmunol.1301721 (2014).

24 Li, Q. et al. Immune response in COVID-19: what is next? Cell Death Differ 29, 1107–1122, doi:10.1038/s41418-022-01015-x (2022).

25 Schmidt, T. et al. Cellular immunity predominates over humoral immunity after homologous and heterologous mRNA and vector-based COVID-19 vaccine regimens in solid organ transplant recipients. Am J Transplant 21, 3990–4002, doi:10.1111/ajt.16818 (2021).

26 Wherry, E. J. & Barouch, D. H. T cell immunity to COVID-19 vaccines. Science 377, 821–822, doi:10.1126/science.add2897 (2022).

27 Keeton, R. et al. Impact of SARS-CoV-2 exposure history on the T cell and IgG response. Cell Rep Med 4, 100898, doi:10.1016/j.xcrm.2022.100898 (2023).

28 Liwsrisakun, C. et al. Neutralizing antibody and T cell responses against SARS-CoV-2 variants of concern following ChAdOx-1 or BNT162b2 boosting in the elderly previously immunized with CoronaVac vaccine. Immun Ageing 19, 24, doi:10.1186/s12979-022-00279-8 (2022).

29 Jo, N. et al. Impaired CD4(+) T cell response in older adults is associated with reduced immunogenicity and reactogenicity of mRNA COVID-19 vaccination. Nature Aging 3, 82-+, doi:10.1038/s43587-022-00343-4 (2023).

30 Britanova, O. V. et al. Age-related decrease in TCR repertoire diversity measured with deep and normalized sequence profiling. J Immunol 192, 2689–2698, doi:10.4049/jimmunol.1302064 (2014).

31 Pereira, B. I. et al. Sestrins induce natural killer function in senescent-like CD8(+) T cells. Nat Immunol 21, 684–694, doi:10.1038/s41590-020-0643-3 (2020).

32 Cheon, I. S. et al. Immune signatures underlying post-acute COVID-19 lung sequelae. Sci Immunol 6, eabk1741, doi:10.1126/sciimmunol.abk1741 (2021).

33 Lo Tartaro, D., et al. Molecular and cellular immune features of aged patients with severe COVID-19 pneumonia. Commun Biol 5, doi: ARTN 590 10.1038/s42003-022-03537-z (2022).

34 Shi, H. B. et al. The inhibition of IL-2/IL-2R gives rise to CD8(+) T cell and lymphocyte decrease through JAK1-STAT5 in critical patients with COVID-19 pneumonia. Cell Death Dis 11, doi: ARTN 429 10.1038/s41419-020-2636-4 (2020).

35 Zhang, Y. et al. Potential contribution of increased soluble IL-2R to lymphopenia in COVID-19 patients. Cell Mol Immunol 17, 878–880, doi:10.1038/s41423-020-0484-x (2020).

36 Zhang, Y. G. et al. Potential contribution of increased soluble IL-2R to lymphopenia in COVID-19 patients. Cellular & Molecular Immunology 17, 878–880, doi:10.1038/s41423-020-0484-x (2020).

37 Wen, W. et al. Immune cell profiling of COVID-19 patients in the recovery stage by single-cell sequencing. Cell Discov 6, 31, doi:10.1038/s41421-020-0168-9 (2020).

38 Toor, S. M., Saleh, R., Sasidharan Nair, V., Taha, R. Z. & Elkord, E. T-cell responses and therapies against SARS-CoV-2 infection. Immunology 162, 30–43, doi:10.1111/imm.13262 (2021).

39 Martinez-Zamudio, R. I. et al. Senescence-associated beta-galactosidase reveals the abundance of senescent CD8+T cells in aging humans. Aging Cell 20, doi: ARTN e13344 10.1111/acel.13344 (2021).

