## supplemental material for "CD4^+^ T cell senescence is associated with reduced reactogenicity in severe/critical COVID-19"

Gating strategy of S1 Panel

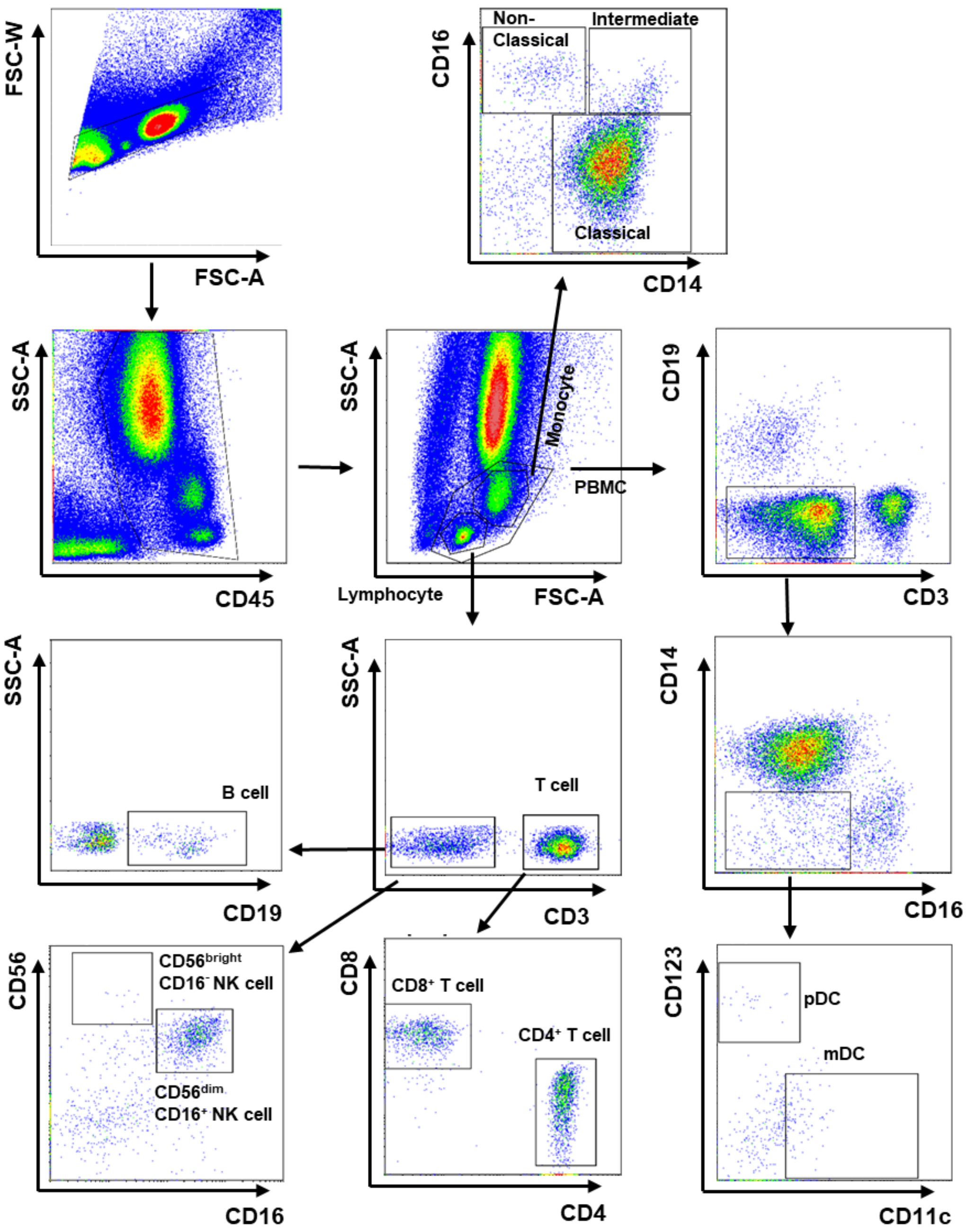

### Gating strategy of S2 Panel

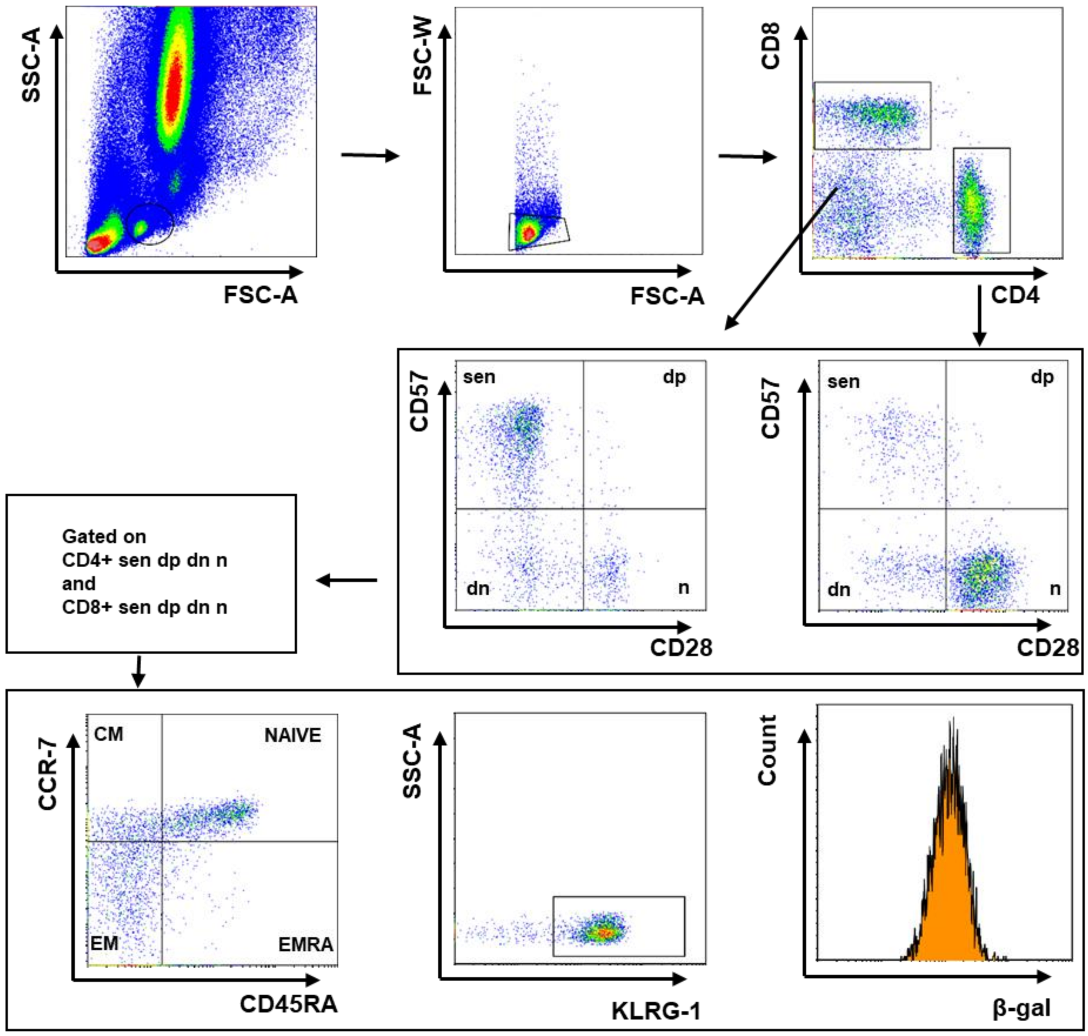

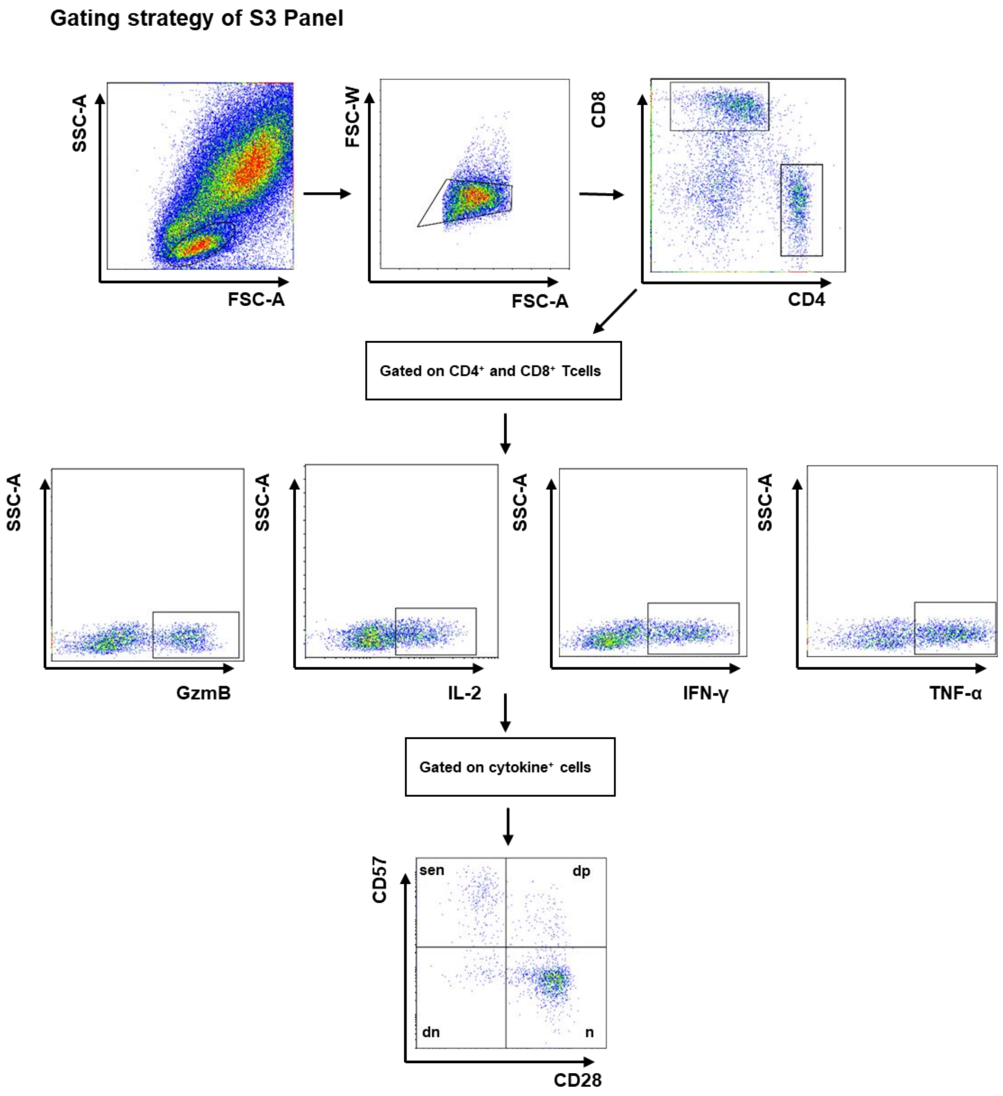


Supplementary Figure 1. The gating strategy of S1, S2 and S3 Panel

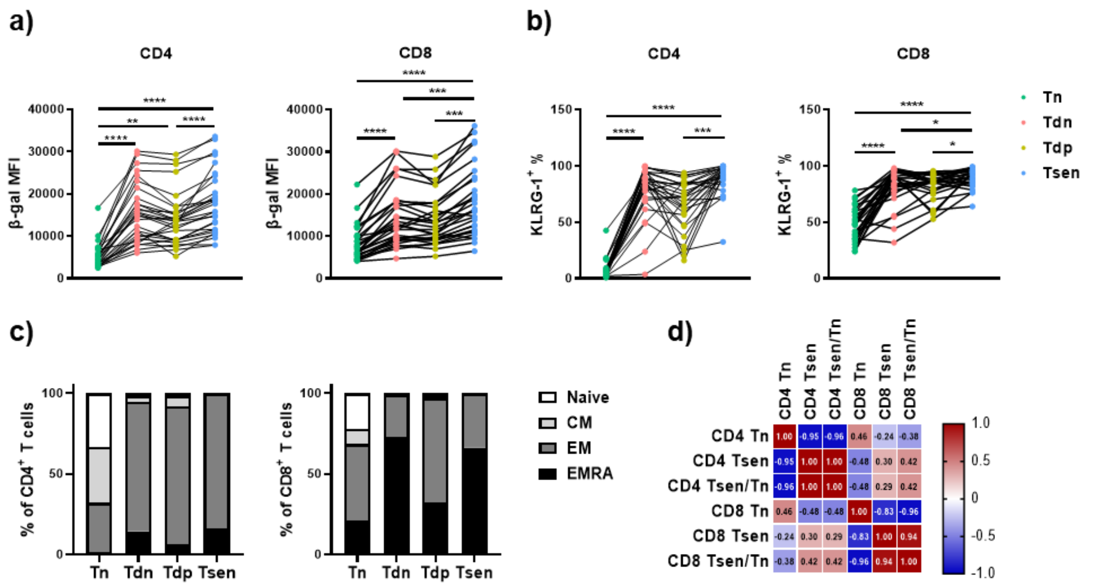


Supplementary Figure 2 Detailed phenotypic presentation of senescent T cells

(a) The mean fluorescence intensity of β-gal in the four subsets (Tn, Tdn, Tdp and Tsen) (n=29) of $CD4^+$ and $CD8^+$ T cells

(b) The percentage of $KLRG1^+$ T cells in the four subsets (Tn, Tdn, Tdp and Tsen) (n=29) of $CD4^+$ and $CD8^+$ T cells

(c) The percentage of naïve, central memory, effect memory and terminal differentiation effect memory T cells in the four subsets (Tn, Tdn, Tdp and Tsen) of $CD4^+$ and $CD8^+$ T cells (n=29)

(d) Correlations between the percentage of 3 subsets (Tn, Tsen, Tsen/Tn) of $CD4^+$ T cells and of $CD8^+$ T cells (n=100)

*, p< 0.05; **, p <0.01; ***, p <0.001; ****, p<0.0001.

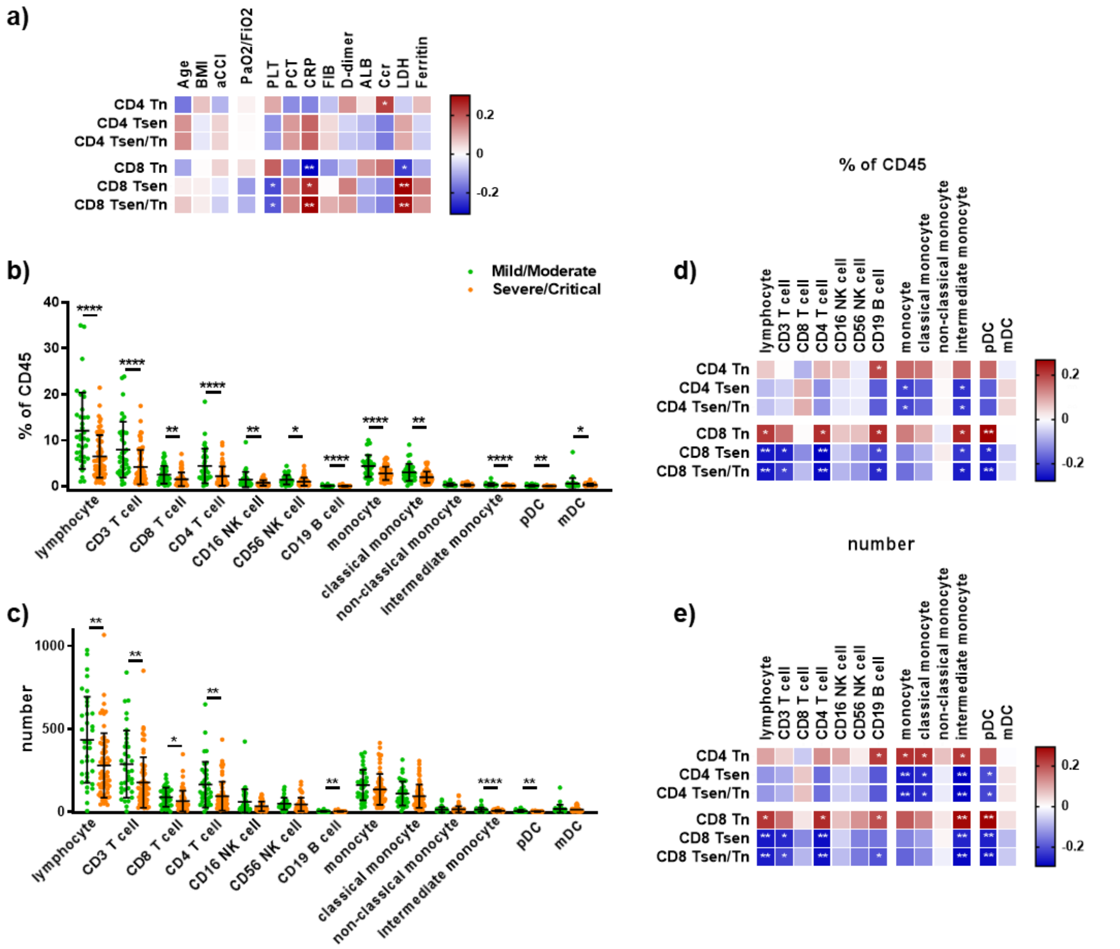


Supplementary Figure 3

(a) Correlations between the 3 subsets (Tn, Tsen, Tsen/Tn) of T cells and clinical phenotypes (n=100)

(b) The percentage of different immune cell subtypes in $CD45^+$ white blood cells of mild/moderate (n=36) or sever/critical (n=64) patients

(c) The number of different immune cell subtypes in mild/moderate (n=36) or mild/moderate (n=64) patients

(d) Correlations between the 3 subsets (Tn, Tsen, Tsen/Tn) of T cells and the percentage of different immune cell subtypes in $CD45^+$ white blood cells (n=100)

(e) Correlations between the 3 subsets (Tn, Tsen, Tsen/Tn) of T cells and the number of different immune cell subtypes (n=100)

*, $p< 0.05$; **, $p <0.01$; ***, $p <0.001$; ****, $p <0.0001$.

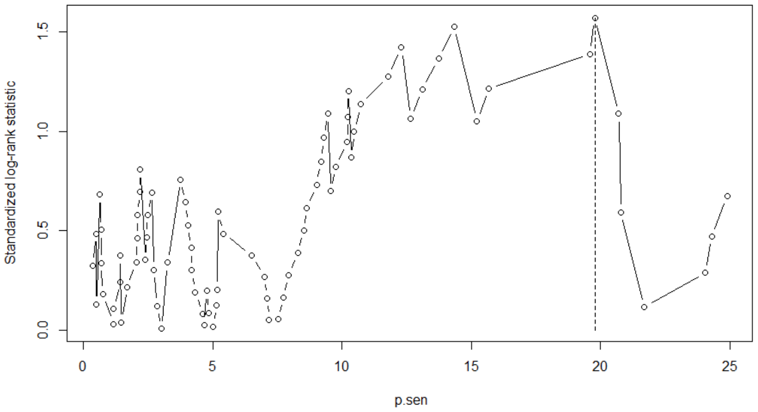


Supplementary Figure 4. Optimization of CD4 Tsen cut-off according to maximization of log-likelihood ratio method

Variation of death rate according to circulating senescent lymphocytes (% CD28-CD57$^{+}$ among CD4$^{+}$ T-cells)

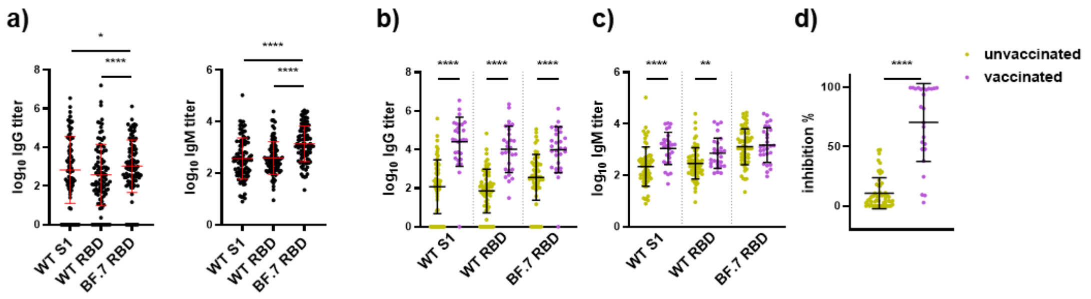


Supplementary Figure 5 The spike specific antibody titers and the inhibition rate of neutralization antibody in COVID-19 patients

(a) The titer of IgG or IgM in plasma against the protein of WT S1, WT RBD and BF.7 RBD (n=97)

(b-c) The titer of IgG or IgM against the protein of WT S1, WT RBD and BF.7 RBD in unvaccinated (n=55) or vaccinated patients (n=26)

(d) The inhibition rate of neutralization antibody in unvaccinated (n=55) or vaccinated patients (n=26)

*, $p< 0.05$; **, $p <0.01$; ***, $p <0.001$ ; ****, $p<0.0001$.

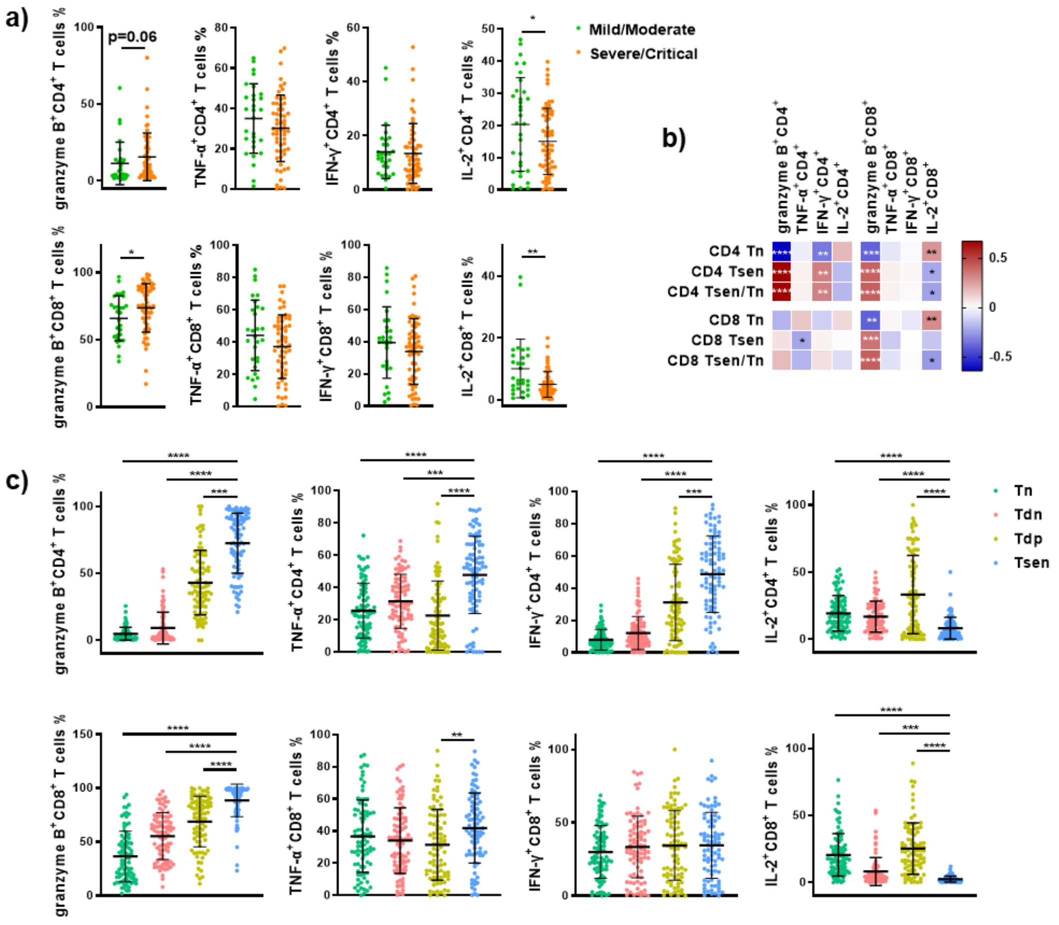


Supplementary Figure 6 The percentage of T cells that release cytokines

(a) The percentage of different cytokines (granzyme B, TNF-α， IFN-γ， IL-2) released $CD4^+$ or $CD8^+$ T cells in mild/moderate (n=29) or sever/critical (n=58) patients

(b) Correlations between the percentage of 3 subsets of T cells (Tn, Tsen, Tsen/Tn) and the percentage of cytokine released T cells (n=87)

(c) The percentage of different cytokines (granzyme B, TNF-α， IFN-γ， IL-2) released subsets (Tn, Tdn, Tdp, Tsen) of $CD4^+$ and $CD8^+$ T cells (n=87)

*, p< 0.05; **, p <0.01; ***, p <0.001 ; ****, p<0.0001.

Supplementary Table 1 Reagent used in this research

| Reagent in Different Panels | Com | Catalog | Clone |
|---|---|---|---|
| **Panel S1** | | | |
| PE anti-human CD4 | Biolegend | 300508 | RPA-T4 |
| PerCP-cy5.5 anti-human CD8 | Biolegend | 344710 | SK1 |
| PE-Cy7 anti-human CD56 | Biolegend | 318318 | HCD56 |
| BV605 anti-human CD16 | Biosciences | 740436 | B73.1 |
| PE-CF594 anti-human CD14 | Biolegend | 325634 | HCD14 |
| APC-Cy7 anti-human CD3 | Biolegend | 300426 | UCHT1 |
| APC anti-human CD123 | Biolegend | 306012 | 6H6 |
| BV421 anti-human CD19 | Biolegend | 302233 | HIB19 |
| BV510 anti-human CD11c | Biolegend | 371513 | S-HCL-3 |
| BV650 anti-human CD45 | Biosciences | 563717 | HI30 |
| **Panel S2** | | | |
| APC-Cy7 anti-human CD4 | Biolegend | 357416 | A161A1 |
| PerCP-cy5.5 anti-human CD8 | Biolegend | 344710 | SK1 |
| PE-Cy7 anti-human CD28 | Biolegend | 302926 | CD28.2 |
| BV421 anti-human CD57 | Biolegend | 359608 | HNK-1 |
| BV650 anti-human CD45RA | Biosciences | 563963 | HI100 |
| PE-CF594 anti-human CCR7 | Biolegend | 353236 | G043H7 |
| APC anti-human KLRG-1 | Biolegend | 367715 | SA231A2 |
| Cellular Senescence Detection Kit-SPiDER-β Gal | | SG03 | |
| **Panel S3** | | | |
| APC-Cy7 anti-human CD4 | Biolegend | 357416 | A161A1 |
| PerCP-cy5.5 anti-human CD8 | Biolegend | 344710 | SK1 |
| PE-Cy7 anti-human CD28 | Biolegend | 302926 | CD28.2 |
| BV421 anti-human CD57 | Biolegend | 359608 | HNK-1 |
| FITC anti-human IFN-γ | Biolegend | 359607 | HNK-1 |
| PE-CF594 anti-human TNF-α | Biolegend | 502946 | Mab11 |
| APC anti-human GranzymeB | Biolegend | 372204 | QA16A02 |
| BV605 anti-human IL-2 | Biolegend | 500331 | MQ1-17H12 |

Supplementary Table 2 Demographics, Characteristics, and Clinical Features of Patients With Coronavirus Disease 2019[a]

| Characteristics | All cases (n=100) | CD4 Tsen low (n= 84) | CD4 Tsen high (n=16) | *P*-value[b] |
|---|---|---|---|---|
| Age, y(n) | 80.10±9.89 | 79.98±9.89 | 80.69±10.22 | 0.797 |
| Sex, male | 64 (64%) | 53 (63.1%) | 11 (68.8%) | 0.666 |
| BMI, kg/m$^2$ | 23.81±3.91 (96) | 23.84±3.97 | 23.69±3.74 | 0.895 |
| <18.5 | 7 (7.3%) | 5 (6.2%) | 2 (13.3%) | 0.578 |
| 18.5-23.9 | 40 (41.7%) | 35 (43.2%) | 5 (33.3%) | |
| 24.0-27.9 | 37 (38.5%) | 30 (37.0%) | 7 (46.7%) | |
| ≥28.0 | 12 (12.5%) | 11 (13.6%) | 1 (6.7%) | |
| Smoking History, yes (n) | 35 (35.0%) | 27 (32.1%) | 8 (50.0%) | 0.170 |
| Any comorbidity | | | | |
| Diabetes | 25 (25%) | 22 (26.2%) | 3 (18.8%) | 0.753 |
| Hypertension | 52 (52.0%) | 40 (47.6%) | 12 (75.0%) | 0.045 |
| Cardiovascular diseases | 24 (24.0%) | 23 (27.4%) | 1 (6.3%) | 0.135 |
| COPD | 11 (11.0%) | 8 (9.5%) | 3 (18.8%) | 0.519 |
| Asthma | 4 (4.0%) | 3 (3.6%) | 1 (6.3%) | 0.508 |
| aCCI | 4.92±1.23 | 4.69±1.25 | 4.85±1.33 | 0.431 |
| Signs and symptoms | | | | |
| Fever | 82 (82.0%) | 68 (81.0%) | 14 (87.5%) | 0.787 |
| Cough | 85 (85.0%) | 70 (83.3%) | 15 (93.8%) | 0.492 |
| Sputum Production | 80 (80.0%) | 66 (78.6%) | 14 (87.5%) | 0.633 |
| Dyspnea | 60 (60.0%) | 52 (61.9%) | 8 (50.0%) | 0.373 |
| Medication | | | | |
| Glucocorticoids | 84 (84.0%) | 71 (81.3%) | 84 (84.0%) | 0.743 |

BMI, body mass index; aCCI, age-adjusted Charlson Comorbidity Index.

a.Continuous variables were presented as mean ± SD (n); categorical variables are shown as n (%). Medication and respiratory support information was recorded during entire hospital stay; other information was recorded at admission.

b.P-values were from t-test for continuous data and from $\chi 2$ test for categorical data

Supplementary Table 3 Laboratory Characteristics on Admission for Severely and Critically Ill Patients with Coronavirus Disease 2019[a]

| Characteristics | All cases (n=100) | CD4 Tsen low (n=84) | CD4 Tsen high (n=16) | *P*-value[b] |
|---|---|---|---|---|
| Blood routine | | | | |
| White blood cell count, $10^9$/L | 7.56±2.9 | 7.42±3.05 | 7.64±2.77 | 0.797 |
| <3.5 | 2（2.0%） | 2（2.4%） | 0（0.0%） | 0.429 |
| 3.5~9.5 | 75（75.0%） | 61（72.6%） | 14（87.5%） | |
| >9.5 | 23（23.0%） | 21（25.0%） | 2（12.5%） | |
| Neutrophil count, $10^9$/L | 6.39±2.77 | 6.42±2.84 | 6.20±2.46 | 0.785 |
| Lymphocyte count, $10^9$/L | 0.78±0.48 | 0.78±0.49 | 0.76±0.38 | 0.686 |
| Platelet count, $10^9$/L | 214.62±81.04 | 216.29±84.38 | 201.75±69.30 | 0.492 |
| Hemoglobin, g/L | 121±28.41 | 121.96±32.37 | 114.69±17.60 | 0.056 |
| Inflammatory markers | | | | |
| Procalcitonin, ng/mL | 0.38±1.10 | 0.29±0.73 | 0.99±2.30 | 0.710 |
| <0.1 | 53（54.1%） | 36（48.09%） | 7（50.0%） | 0.883 |
| 01~0.3 | 30（30.6%） | 26（34.7%） | 4（28.6%） | |
| >0.3 | 15（15.3%） | 13（17.3%） | 3（21.4%） | |
| C-reactive protein, mg/L | 24.76±40.66 | 25.14±37.66 | 44.04±63.40 | 0.037 |
| ≤8 | 41（42.7%） | 36（44.4%） | 45（55.6%） | 0.424 |
| >8 | 55（57.3%） | 5（33.3%） | 10（66.7%） | |
| Coagulation function | | | | |
| D-dimer, ug/mL | 2.77±5.00 | 2.86±4.77 | 1.08±1.17 | 0.387 |
| ≤age/100 | 47（49.0%） | 38（47.5%） | 42（52.5%） | 0.523 |
| > age/100 | 49（51.0%） | 9（56.3%） | 7（43.8%） | |
| Serum biochemical indicators | | | | |
| Serum albumin level, g/L | 31.89±4.87 | 31.89±4.05 | 31.19±4.32 | 0.531 |
| Creatinine, μmol/L | 98.34±92.85 | 96.75±71.64 | 99.23±103.42 | 0.413 |
| Serum urea nitrogen, mmol/L | 9.96±8.53 | 10.29±9.20 | 8.48±4.52 | 0.472 |
| Total bilirubin, μmol/L | 12.02±5.66 | 12.52±6.28 | 9.67±2.96 | 0.227 |
| Alanine Aminotransferase,U/L | 37.49±38.22 | 38.50±38.75 | 31.85±19.36 | 0.525 |
| Aspartate Aminotransferase, U/L | 42.47±35.11 | 42.86±27.99 | 41.31±19.70 | 0.821 |
| Creatine kinase, U/L | 109.06±182.15 | 111.80±192.01 | 94.07±117.18 | 0.151 |
| Creatine kinase-MB, U/L | 15.56±31.62 | 17.11±34.32 | 7.50±3.03 | 0.024 |

a. Continuous variables were presented as median (interquartile range); categorical variables are shown as n (%).

b. P-values were from t-test for normally distributed continuous data and from Mann-Whitney U test for abnormally distributed continuous data.P-values were from $\chi2$ test for categorical data.

Supplementary Table 4 Plasma soluble factors in COVID-19 patients

| Cytokines | Mild/Moderate (n=19) | Severe/Critical (n=34) | *P*-value[b-0] |
|---|---|---|---|
| IL-2 | 63.46±15.86 | 58.34±9.993 | 0.3852 |
| IL4 | 39.51±30.01 | 12.17±4.094 | 0.2385 |
| Il-10 | 5.809±2.370 | 12.39±2.627 | 0.1013 |
| IL-6 | 22.02±9.565 | 71.54±23.36 | 0.1300 |
| Il-17a | 5.221±1.582 | 3.519±0.8046 | 0.2913 |
| TNF-α | 14.86±7.479 | 14.28±3.242 | 0.9345 |
| sFas | 544.0±70.80 | 701.1±63.76 | 0.1240 |
| sFasL | 18.27±3.782 | 13.22±1.946 | 0.1940 |
| IFN-γ | 123.9±48.29 | 180.7±49.86 | 0.4578 |
| GranzymeA | 171.2±95.14 | 102.5±19.70 | 0.3659 |
| GranzymeB | 467.8±133.9 | 473.7±101.8 | 0.9723 |
| Perforin | 1050±106.2 | 914.5±72.63 | 0.2859 |
| Granulysin | 2399±143.8 | 2423±195.5 | 0.9341 |
| IL8 | 11.21±4.082 | 16.36±5.718 | 0.5351 |
| IP10 | 522.2±182.0 | 1228±255.5 | 0.0612 |
| Eotaxin | 79.40±10.17 | 75.35±6.513 | 0.7271 |
| TARC | 40.55±7.107 | 38.25±6.708 | 0.8266 |
| MCP1 | 164.4±37.54 | 278.2±68.7 | 0.2434 |
| RANTES | 953.0±99.30 | 931.9±119.4 | 0.9056 |
| MIP1α | 7.077±3.271 | 9.218±7.993 | 0.8466 |
| MIG | 492.9±111.6 | 758.0±141.0 | 0.2054 |
| ENA78 | 19.69±4.668 | 23.79±5.070 | 0.5937 |
| MIP3α | 10.84±3.767 | 10.23±2.242 | 0.8816 |
| GROα | 23.08±6.923 | 17.57±3.826 | 0.4516 |
| ITAC | 30.68±5.317 | 61.00±14.72 | 0.1389 |
| MIP1β | 2.773±0.5034 | 4.034±1.288 | 0.4791 |
| TSLP | 7.891±5.190 | 9.460±4.710 | 0.8331 |
| IL1α | 37.59±13.15 | 26.94±8.934 | 0.4941 |
| IL1β | 11.53±3.404 | 12.43±2.684 | 0.8386 |
| GM-CSF | 3.230±2.045 | 6.398±2.057 | 0.3198 |
| IFNα2 | 5.942±1.260 | 4.956±0.9000 | 0.5221 |
| IL23 | 53.35±13.81 | 46.99±6.493 | 0.6383 |
| IL12p40 | 85.19±18.50 | 75.34±17.31 | 0.7169 |
| IL12p70 | 2.748±1.025 | 4.177±0.7668 | 0.2697 |
| IL15 | 152.8±39.19 | 115.3±22.73 | 0.3779 |
| IL18 | 89.88±32.42 | 137.735.91 | 0.3796 |
| IL11 | 108.9±33.62 | 140.5±28.84 | 0.4965 |
| IL27 | 73.97±16.57 | 66.37±18.42 | 0.7843 |
| IL33 | 63.12±27.80 | 43.12±13.72 | 0.4729 |
| IL5 | 3.911±1.476 | 7.316±1.396 | 0.1231 |
| IL13 | 7.809±4.382 | 16.50±4.347 | 0.1993 |
| IL9 | 23.78±15.36 | 11.18±1.710 | 0.2836 |
| IL17F | 5.537±4.154 | 3.267±0.609 | 0.4798 |

| | | | |
|---|---|---|---|
| IL22 | 6.358±2.188 | 4.644±0.8429 | 0.3904 |

a. Different cytokine concentrations (pg/mL) of mild/moderate or severe/critical patients in plasma were presented as mean ± SEM

b. P-values were from t-test for normally distributed continuous data and from Mann-Whitney U test for abnormally distributed continuous data.
